# An agent-based model of insect resistance management and mitigation for Bt maize: A social science perspective

**DOI:** 10.1101/732776

**Authors:** Yuji Saikai, Paul D. Mitchell, Terrance M. Hurley

## Abstract

Managing and mitigating agricultural pest resistance to control technologies is a complex system in which biological and social factors spatially and dynamically interact. We build a spatially explicit population genetics model for the evolution of pest resistance to Bt toxins by the insect *Ostrinia nubilalis* and an agent-based model of Bt maize adoption, emphasizing the importance of social factors. The farmer adoption model for Bt maize weighed both individual profitability and adoption decisions of neighboring farmers to mimic the effects of economic incentives and social networks. The model was calibrated using aggregate adoption data for Wisconsin. Simulation experiments with the model provide insights into mitigation policies for a high-dose Bt maize technology once resistance emerges in a pest population. Mitigation policies evaluated include increased refuge requirements for all farms, localized bans on Bt maize where resistance develops, areawide applications of insecticidal sprays on resistant populations, and taxes on Bt maize seed for all farms. Evaluation metrics include resistance allele frequency, pest population density, farmer adoption of Bt maize and economic surplus generated by Bt maize.

Based on economic surplus, the results suggest that refuge requirements should remain the foundation of resistance management and mitigation for high-dose Bt maize technologies. For shorter planning horizons (< 16 years), resistance mitigation strategies did not improve economic surplus from Bt maize. Social networks accelerated the emergence of resistance, making the optimal policy intervention for longer planning horizons rely more on increased refuge requirements and less on insecticidal sprays targeting resistant pest populations. Overall, the importance social factors play in these results implies more social science research, including agent-based models, would contribute to developing better policies to address the evolution of pest resistance.

**Author Summary:** Bt maize has been a valuable technology used by farmers for more than two decades to control pest damage to crops. Using Bt maize, however, leads to pest populations evolving resistance to Bt toxins so that benefits decrease. As a result, managing and mitigating resistance has been a serious concern for policymakers balancing the current and future benefits for many stakeholders. While the evolution of insect resistance is a biological phenomenon, human activities also play key roles in agricultural landscapes with active pest management, yet social science research on resistance management and mitigation policies has generally lagged biological research. Hence, to evaluate policy options for resistance mitigation for this complex biological and social system, we build an agent-based model that integrates key social factors into insect ecology in a spatially and dynamically explicit way. We demonstrate the significance of social factors, particularly social networks. Based on an economic surplus criterion, our results suggest that refuge requirements should remain the foundation of resistance mitigation policies for high-dose Bt technologies, rather than localized bans, areawide insecticide sprays, or taxes on Bt maize seed.

## Introduction

Globally, farmers have planted more than 2.3 billion hectares of genetically engineered crops since their commercial introduction in 1996, including a new maximum of 190 million hectares in 2017 [1]. Focusing on maize (*Zea mays*), the world’s leading grain crop with annual production exceeding a billion metric tons, the United States, Brazil and Argentina together produced almost half of the world’s supply in 2017 [2]. Bt maize – maize genetically engineered to produce *Bacillus thuringiensis* (Bt) toxins in plant tissues for insect control – accounted for more than 80% of the maize planted in each of these three nations in 2017 [1]. After more than two decades of commercial use of genetically engineered crops, insect resistance to Bt toxins continues to be a major concern around the world [3]. A high- dose/refuge resistance management strategy continues to be the primary policy in multiple nations for delaying resistance to these Bt toxins [4–6]. Nevertheless, field-evolved resistance to some of these Bt toxins has been documented for populations of western corn rootworm (*Diabrotica virgifera virgifera*) in the United States and various lepidopteran species in multiple locations [7].

The commercialization of Bt crops has generated a variety of research, including bioeconomic models that integrate population genetics and pest ecology with farmer economic returns [8–10]. Though these models contributed to the development of insect resistance management policies, little other work exists on the role of social factors in the evolution of insect resistance to commercialized toxins. Insect resistance to these toxins evolves in response to human management activities, activities driven by a variety of social factors that include not only economic considerations, but also sociological, psychological, cultural, historical and political considerations [11]. As a result, examining genetic and ecological processes in isolation from these broader social factors driving human behavior potentially misses key determinants of the evolution of insect resistance. Hence, a broader, complex systems model of insect resistance management that incorporates both biological and social processes can potentially provide new insights [12].

In the United States (US), the Environmental Protection Agency (EPA) required companies commercializing Bt crops to develop resistance mitigation plans as a condition for product registration [13]. Once a resistant population has been officially documented according to the EPA process, these resistance mitigation plans generally restrict the availability of the technology (Bt seed) in and around the region where the resistant population emerges. Though resistant insect populations and field failures in the US have been documented in the scientific literature [7, 14], the official EPA criteria have yet to trigger implementation of these mitigation plans for any pest. Instead, the EPA has required a more generalized response by Bt crop registrants [15]. Interestingly, little research exists that evaluates and compares the mitigation plans that have been filed or other mitigation policies, particularly from an economic perspective. Given the length of time that Bt crops have been in use in the US and elsewhere, insect resistance is likely to become an increasing problem, making more research on mitigation responses and strategies especially timely.

This paper has two goals. First, we develop an agent-based model of insect resistance to Bt maize that incorporates farmer adoption behavior. We then use the model to compare different mitigation policies in order to inform policymakers and other stakeholders of the types of programs that are likely to generate the largest economic benefits for society. Second, focusing specifically on the impact of social networks on farmer adoption behavior, we show that social factors can also play a key role in the evolution of insect resistance to Bt toxins in agricultural cropping systems.

Agent-based modeling has become more widely-used for studying complex systems and emergent behavior, including socio-ecological modeling of insect resistance management [16, 17]. In agent-based models, an observed macroscopic phenomenon emerges as a result of interaction among heterogeneous agents in a dynamically evolving environment. Agents typically follow simple decision rules and influence each other either directly or indirectly through the environment, which itself evolves according to its own rules and agent actions. Because the processes being explicitly modeled are complex, researchers use computer simulations to examine outcomes over a wide range of parameter values. In short, agent-based models are laboratory experiments conducted *in silico* [17, 18]. Despite the remaining challenges to overcome, such as ad hoc assumptions and lack of relevant data for validation [19–21], agent-based modeling can provide insights into complex systems that would be difficult to study otherwise. Given the merits, applications of agent-based models to pest and resistance management in agricultural systems have been developed [22–24].

Although agent-based models can integrate many factors, they still face the fundamental tradeoff in modeling: fidelity to the phenomenon being examined and abstraction for ease of analysis and interpretation [17]. This paper focuses on deriving new insights into policy options for mitigating insect resistance once it has evolved, and emphasizes the significance of social factors for questions relevant to policymakers [22]. As a result, social components are richer than existing models that use individual-based modeling to incorporate social factors [25], while the biological aspects of the model are simpler than other models focusing on biological processes [26–28].

We extend existing work [25] on insect resistance management for Bt crops by more fully leveraging the power of agent-based modeling. First, we explicitly model the local influence that neighbors have on farmers through social networks as they make decisions regarding adoption of Bt maize, creating a hybrid decision process that mixes both individual profit considerations and a desire to mimic neighbors. Second, we allow the additional cost of planting Bt seed to vary over time, because this cost influences adoption decisions and companies have reduced the cost of single-toxin Bt seed to encourage farmers to continue to plant Bt maize in the face of pest population suppression [29]. With this pricing flexibility, we calibrate the farmer decision model using historical data that reflects these decreasing prices, and then can examine the impact of a tax on Bt seed as a policy option for mitigating resistance.

For this analysis, we parameterize a bioeconomic model of maize production with the option to use high-dose Bt maize to manage European corn borer (*Ostrinia nubilalis*). We calibrate the Bt maize adoption model using aggregate historical adoption data for farmers in the state of Wisconsin. Through the calibration process, we emphasize the significant role that a social factor – the local influence of social networks on Bt maize adoption [30] – can play in the evolution of insect resistance. Using the calibrated model, we then simulate a number of mitigation policies implemented either over the entire landscape or around the areas where resistance develops. In particular, we consider combinations of an increased refuge requirement and a tax on the sale of Bt seed for all farms, and a ban on the use of Bt maize and areawide use of an additional insecticide to control the pest in the area around where resistance emerges. To assess the relative performance of each policy, we use economic surplus as a monetary measure of the social value generated by the use of Bt maize and conduct sensitivity analysis of key parameters to explore the robustness of model results.

## Results

### Baseline Results

Running the calibrated model 1,000 times with different random seeds and averaging over these iterations gave baseline results for the insect population, the Bt seed adoption rate, and the resistance (R) allele frequency at the landscape level. In the model, periods 0 to 10 were an initialization phase, periods 11 to 32 were a calibration phase corresponding to years 1996 to 2017, and periods 33 to 60 were projections (see Model). The baseline model captured the aggregate Bt adoption rate of Wisconsin farmers by calibrating two parameters that determined Bt maize adoption – farmer responsiveness to profit incentives and the farmer tendency to mimic the adoption decisions of neighbors due to social network effects. The calibrated model reproduced the previously noted oscillation of the European corn borer population before the advent of Bt maize [31], and the documented suppression of the pest population due to the widespread farmer adoption of Bt maize in Wisconsin and other states [32]. As expected, the calibrated model projected a surge in the R-allele frequency as the insect resistance developed, resulting in the eventual recovery of the pest population. Baseline results suggested that period 33 was the beginning of a significant increase in the R-allele frequency. In period 33, the R-allele frequency was 4.1%, but rose quickly, exceeding 10% in period 36, 20% in period 38, 30% in period 39, 40% in period 40 and 50% in period 41. The pest population did not recover until later, with the average density not exceeding 0.5 larvae per plant until period 50.

### Policy Experiments

We simulated policies to mitigate resistance to the Bt toxin once it emerged. Refuge requirements have been the lynch pin of resistance management, and so the mitigation policies we examined began with increasing refuge requirements. In addition, building on the model’s capacity for capturing the complexity from the interaction of biological and social factors, we experimented with combinations of three other types of mitigation policies: localized bans on the use of Bt maize around areas where resistance emerged, areawide applications of other insecticides to control the pest around areas where resistance emerged, and a uniform tax on the sale of Bt seed for all farmers buying it. Refuge policies and localized bans directly regulate the use of Bt maize, areawide spray policies directly manage resistant pest populations, and the Bt seed tax adjusts farmer incentives to use Bt maize. The simulation of resistance mitigation policies was a combination of different assumptions for these four policy parameters: the refuge requirement, localized bans, areawide management, and a Bt seed tax.

The simulated landscape consisted of a grid of fields, with 44% of fields assigned randomly to maize production initially and the remainder to non-maize. Fields remained in their initial allocation throughout a simulation, but were reassigned for each simulation. During a simulation, insect resistance was declared when the R-allele frequency exceeded 50% in the pest population in a field after Bt toxin mortality and before pest dispersal occurred. The 50% threshold was chosen because at this level the landscape-average pest population began to increase (Fig 1), implying that higher population densities were occurring in some fields due to resistance. For resistance mitigation, the refuge requirement was increased from the baseline of 5% to either 20% or 50% for all farmers on the landscape planting Bt maize, with complete compliance achieved using seed mixtures. The localized ban was imposed only on farms within a radius *r* of any field where resistance was declared, again with complete compliance assumed. We considered two radii: once and twice the distance of adult dispersal from the natal field (*r* = 1×dispersal, *r* = 2×dispersal). Conceptually, this ban was a 100% refuge requirement applied locally and dynamically imposed and lifted according to the situation in the previous period. For areawide management, a non-Bt insecticide was applied in the period when resistance was declared, either covering only the field of resistance or all maize fields in a neighborhood around the field within the distance of adult dispersal (*r* = 0×dispersal, *r* = 1×dispersal). We assumed 100% compliance with the insecticide application for all fields within this area and that the application reduced the pest population by 80% after Bt toxin mortality and increased farmer costs by $33.51 ha^-1^. This cost was based on published survey averages for active ingredient and application costs and adjusted for inflation to 2017 equivalents [33, 34]. Finally, the tax policy increased the Bt seed cost by 25% or 50% for all farmers on the landscape for all periods after resistance was declared. In brief, each policy parameter had the following three levels: refuge requirement (5%, 20%, 50%), localized ban (none, *r* = 1×dispersal, *r* = 2×dispersal), areawide spray (none, *r* = 0×dispersal, *r* = 1×dispersal), and Bt seed tax (0%, 25%, 50%). Three levels for each of these four policy parameters created 3^4^ = 81 mitigation policy combinations to simulate.

**Fig 1.**
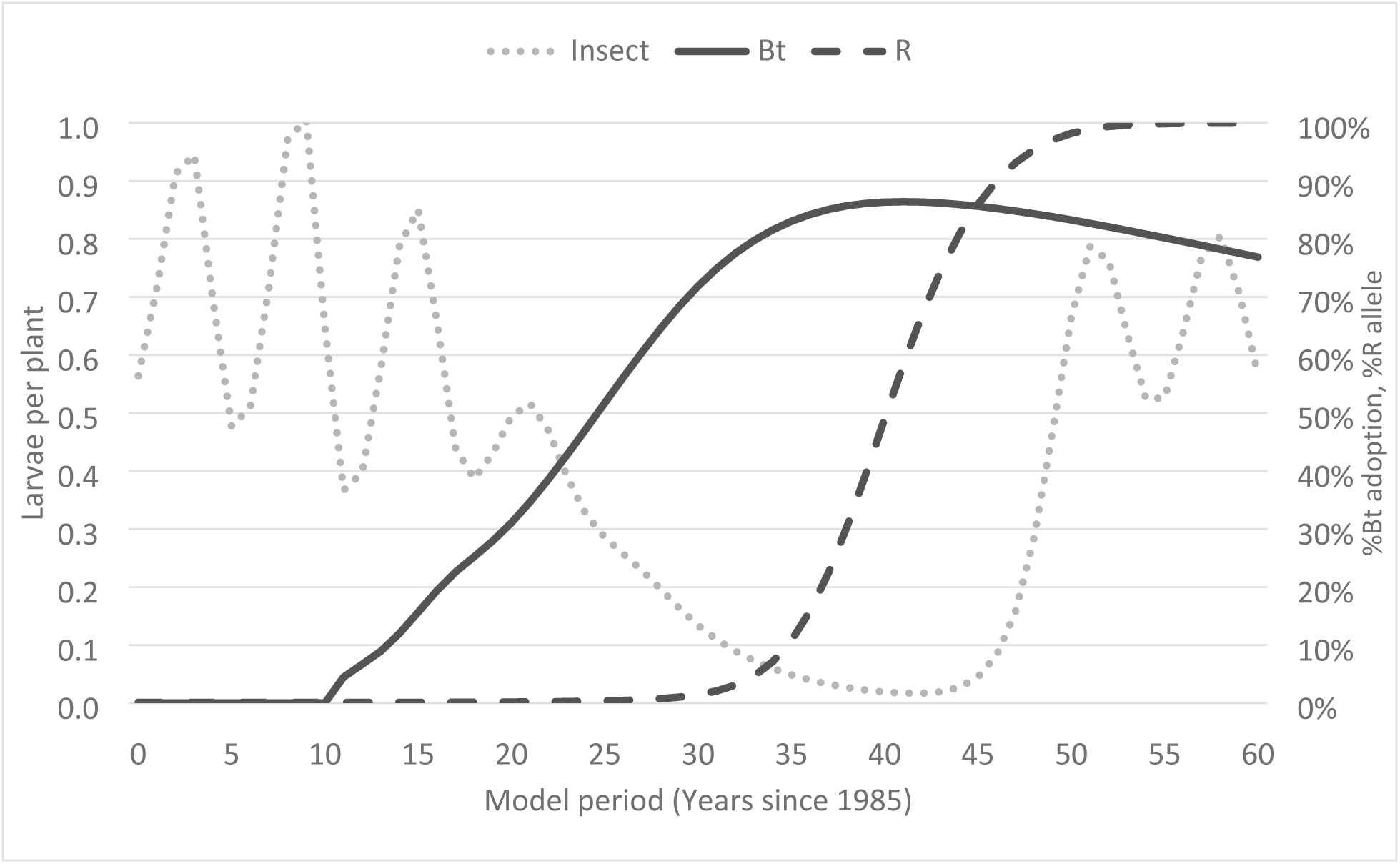
Baseline results from the calibrated model. It contains the insect population density (Insect), Bt seed adoption rate (Bt), and the resistance allele frequency (R) (results for each period are averages over 1,000 simulations).

The calibrated model was run 1,000 times for each policy and, just as for the baseline, the following three results variables were averaged over all 1,000 iterations for each period: aggregate farmer adoption of Bt maize, population-level R-allele frequency, and average pest population density for the landscape. In addition, as a performance metric to compare each policy, we approximated economic surplus each period as the sum of farmer profits and the technology fees collected by the seed company, divided by the total number of farmers.

Costs for spraying insecticides were subtracted from farmer profits for those making applications, while collected taxes were subtracted from farmer profits, but added to the economic surplus (i.e., the tax was a surplus transfer, not a surplus loss). To simplify the analysis, we did not discount future surpluses. Each policy scenario began after the calibration phase (i.e., at period 33), and the cumulative surplus was evaluated for each length of planning horizon ranging from 1 to 25 years (i.e., periods 33 to 57).

To build intuition about the nature of each policy treatment (i.e. refuge, tax, spray, and ban), we first report results for each policy individually (not combinations of policies) by plotting the dynamics for Bt adoption, the R-allele frequency, and the pest population density (Fig 2– Fig 4). In Fig 2 (Bt adoption), results for the ban policy (Ban 1x) are plotted with a separate vertical axis due to its qualitatively different and much stronger effect than for the other policies. Also, results for the spray policy with *r* = 1×dispersal (Spray 1x) and the ban policy with *r* = 2×dispersal (Ban 2x) are omitted as they were very similar to those with smaller radii. In total, Fig 2 plots the Bt adoption rate against the planning horizon for the following policies: baseline (Baseline), 20% refuge (20% Refuge), 50% refuge (50% Refuge), 25% seed tax (25% Tax), 50% seed tax (50% Tax), areawide spray in the field with resistance (Spray 0x) and a localized ban on Bt seed within one pest dispersal radius of the field with resistance (Ban 1x). Consistent with Fig 1, the baseline policy showed a continuing increase in Bt maize adoption from planning horizon year 0 (period 32 in Fig 1), with a peak of almost 86.5% in planning horizon year 9 (period 41 in Fig 1). All policies showed this same general trend (with one exception), but with a lower adoption peak occurring sooner for the seed tax policies, a higher adoption peak occurring later for the increased refuge policies (especially for 50% refuge), and a slightly higher and later peak occurring for the areawide spray policies. The one exception were localized bans the sale of Bt seed, for which implementation caused a rapid decline in the use of Bt maize, with almost complete dis-adoption by the end of the simulation in horizon period 25.

**Fig 2.**
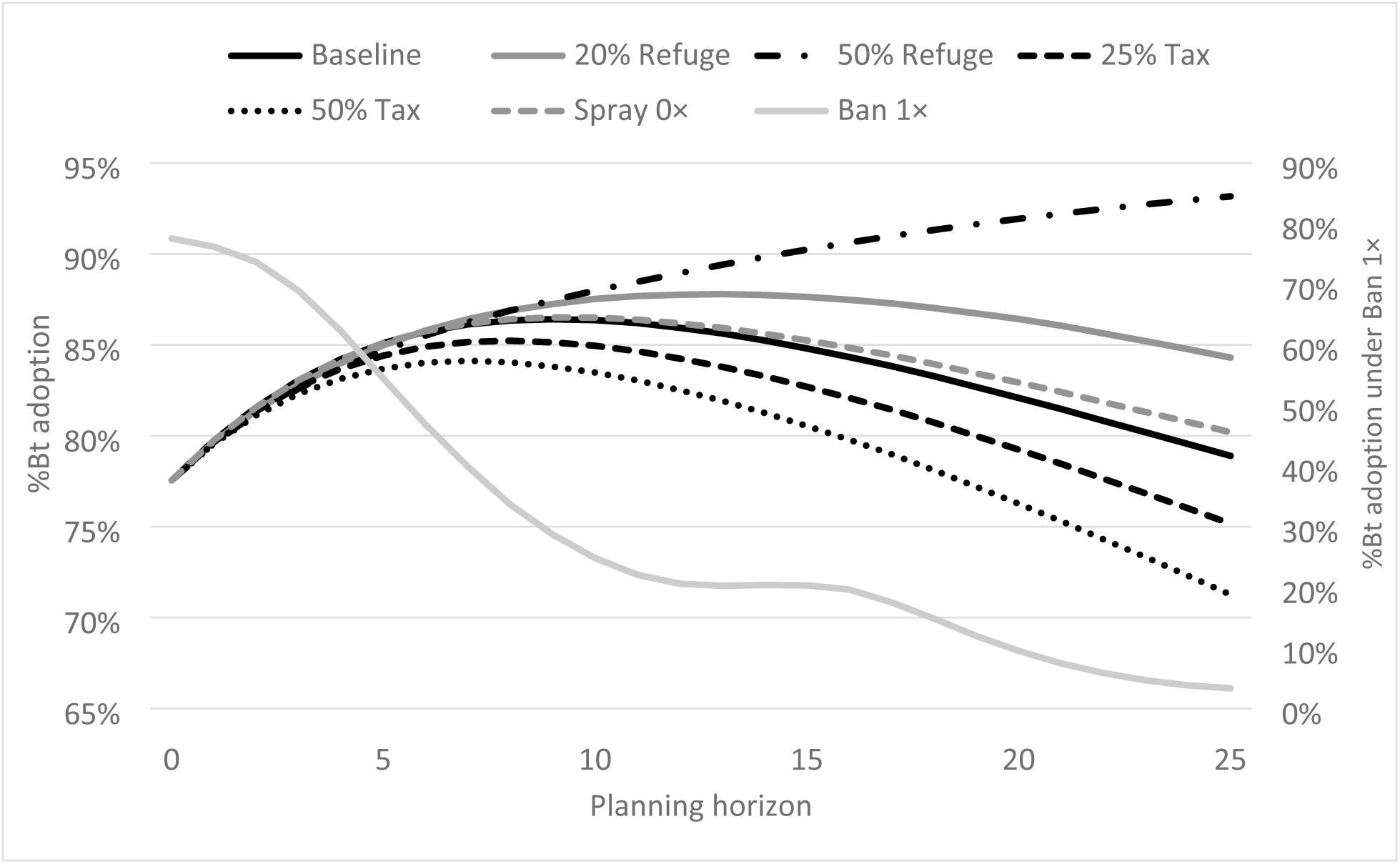
Bt adoption rate under single policies plotted against the planning horizon. The results for each period are averages over 1,000 simulations.

In Fig 3 (R-allele frequency), results for both tax policies are omitted as they were almost identical to the results for the baseline, suggesting low policy efficacy. This result was surprising because Bt adoption differs noticeably for these policies (Fig 2). All mitigation policies plotted in Fig 3 slowed the development of resistance compared to the baseline. The most effective mitigation policies were the 50% refuge for all farms (50% Refuge) and a localized ban on and around fields with resistance (Ban 1x), both of which kept the R-allele frequency below 20% for more than 20 years. By horizon period 25, however, the 50% refuge policy showed a rapid increase in the R-allele frequency, suggesting its failure, while the ban policy kept the frequency below 20%, suggesting that it was the most effective policy for mitigating resistance over the long-run (>25 years). The 20% refuge for all farms (20% Refuge) effectively mitigated the resistance for about 10 years, and then the R-allele frequency began a rapid increase, reaching the baseline level by horizon period 25. The spray policies (Spray 0x) were not particularly effective for mitigating resistance, showing a steady increase in the R-allele frequency, though slower than for the baseline and tax policies.

**Fig 3.**
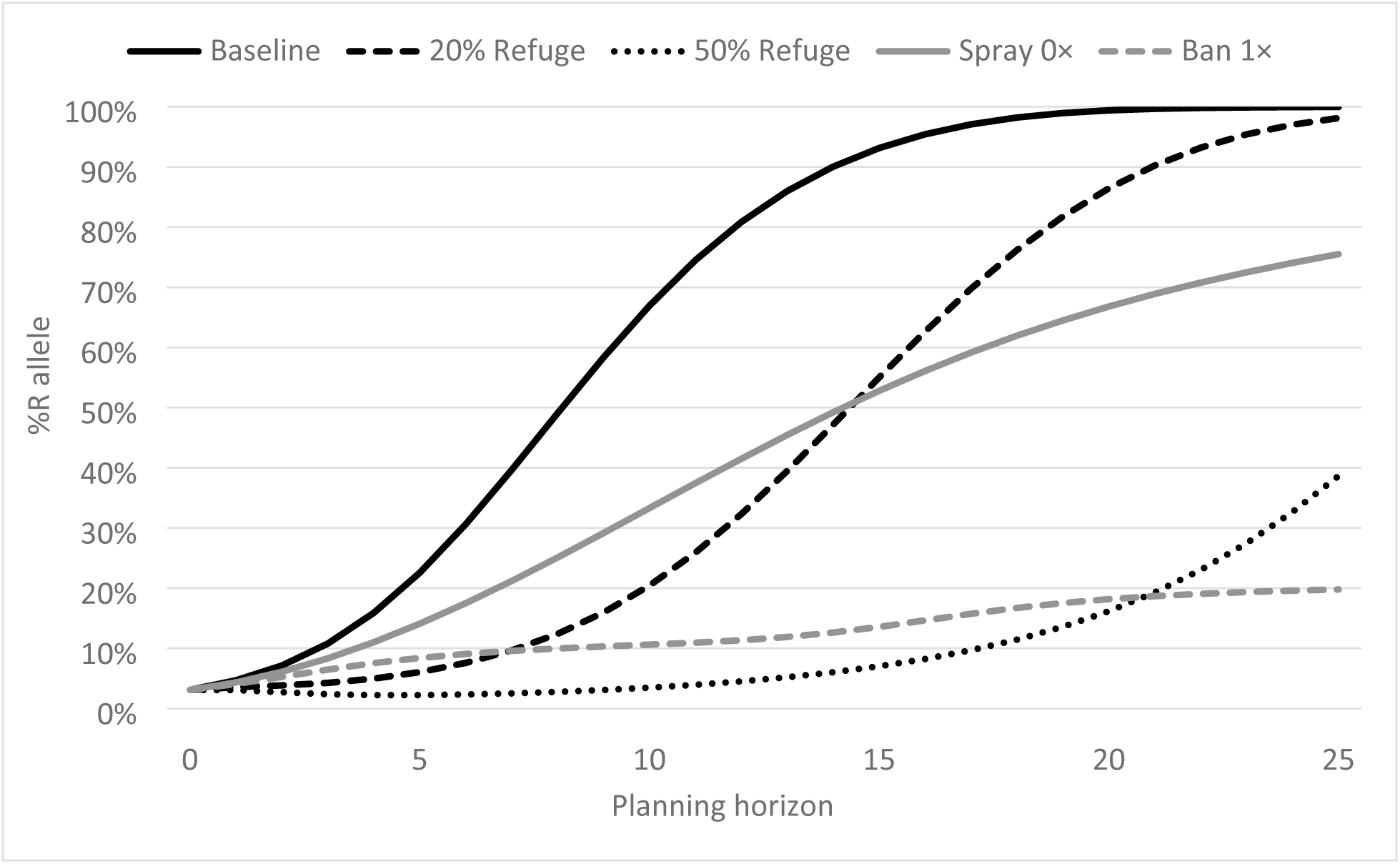
R-allele frequency under single policies plotted against the planning horizon. The results for each period are averages over 1,000 simulations.

**Fig 4.**
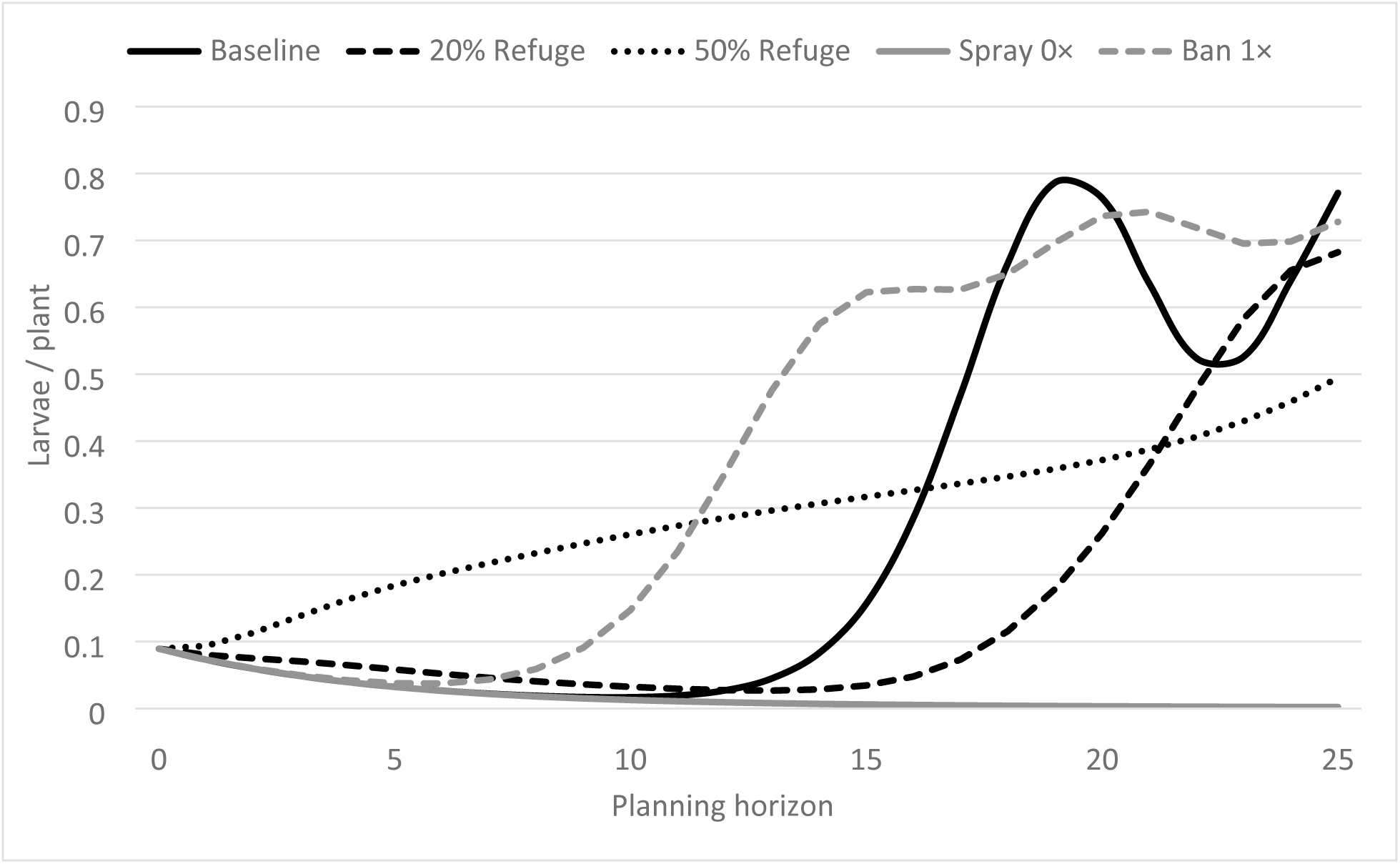
Insect population density under simple policies plotted against the planning horizon. The results for each period are averages over 1,000 simulations.

In Fig 4 (pest population density), results for both tax policies are again omitted as they were almost identical to results for the baseline. The areawide spray policy (Spray 0x) kept the pest population density low over all 25 years, even with a radius = 0, due to the efficacy of the insecticide spray. The baseline with no intervention to mitigate resistance kept the pest population density low for about 15 years, and then the population increased and began to oscillate as expected. Surprisingly, the refuge policies showed distinctly different patterns over the 25 years. The 20% refuge policy (20% Refuge) kept the pest population low for about 20 years (about 5 years longer than the baseline), while the 50% refuge (50% Refuge) showed a long slowly increasing pest population density over all 25 years, exceeding the baseline in year 17 and the 20% refuge policy in year 23. Interestingly, the ban policy (Ban 1x) only kept the pest population low for about 10 years (about 5 years longer than the baseline).

These results showed the tradeoffs inherent in the mitigation of resistance. For example, the 50% refuge and ban policies were both the most effective at reducing the frequency of resistance alleles (Fig 3), but came at the cost of reduced adoption of Bt maize (Fig 2) and higher average pest populations (Fig 4), both implying lower benefits. Hence, we used economic surplus as a measure that integrates across costs and benefits in order to compare mitigation policies and to develop recommendations.

Fig 5 plots average annual economic surplus against the planning horizon, again omitting results for the tax policies as they were almost identical to baseline results. Each point on the curves in Fig 5 is an annualized average of the accumulated surplus over the corresponding planning horizon, i.e., the sum of landscape surplus over the planning horizon, divided by the number of years in the planning horizon and the maize planted area. As seen in Fig 5, the baseline (5% refuge, no localized ban, no areawide spray, no Bt seed tax) generated the greatest average annual surplus for all planning horizons up to 15 years. This result occurred because the surplus measure was cumulative, and the baseline policy accumulated more surplus during the early years than the other policies. However, for planning horizons of 16 or more years, the optimal mitigation policy increased the refuge requirement from 5% to 20% for all farms, but did not impose a localized ban, an areawide spray, or Bt seed tax. The areawide spray was suboptimal due to the additional costs incurred by farmers, while the ban policy was sub-optimal due to the loss of Bt maize benefits for farmers and the lost revenue for the seed company. Interestingly, the 50% refuge policy generated the lowest economic surplus – though it was one of the most effective mitigation policies, its cost in terms of lost benefits to farmers was too high. Recall that results for the two omitted tax policies (25% Tax, 50% Tax) were almost identical to the baseline.

**Fig 5.**
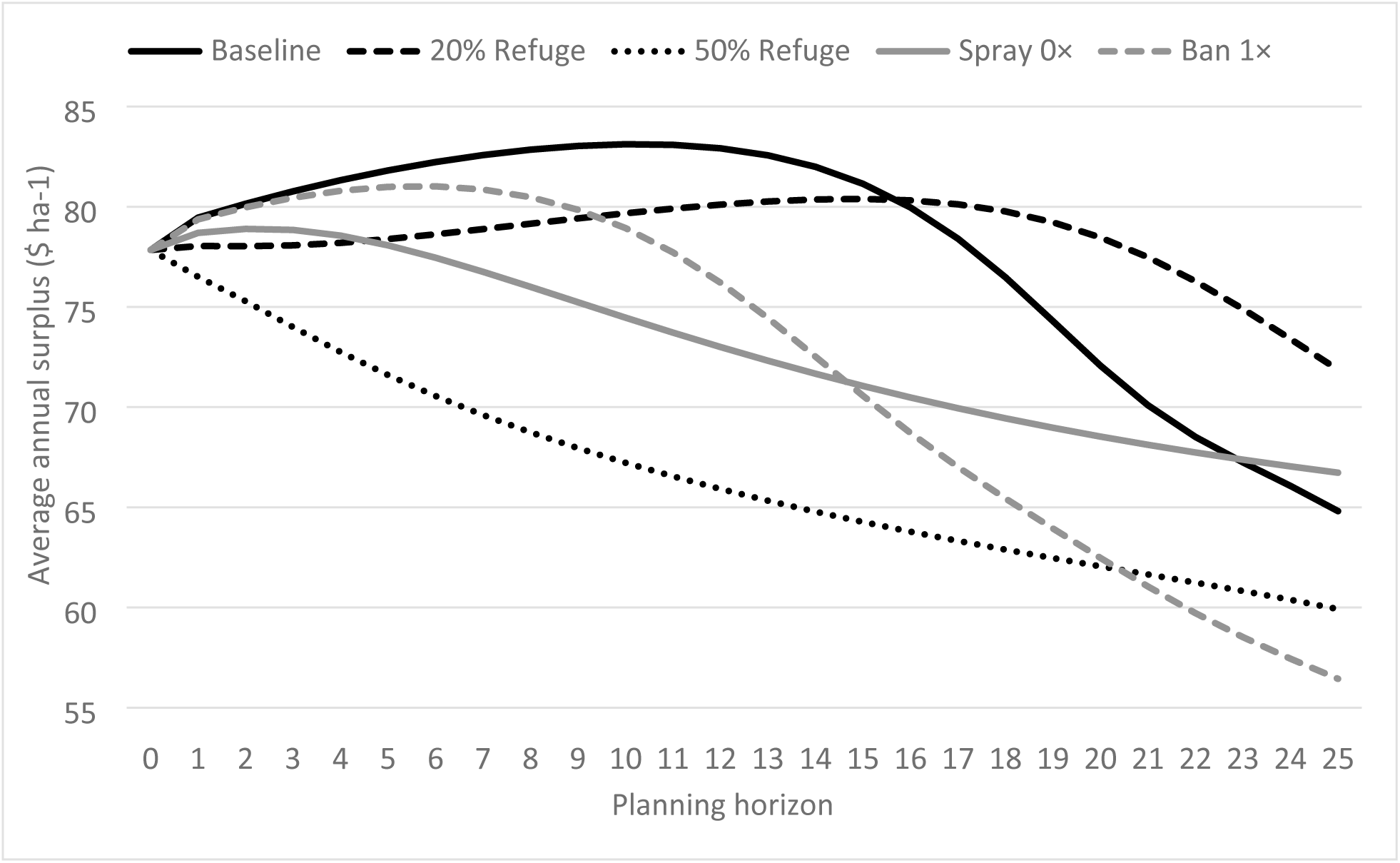
Average annual surplus for different mitigation policies plotted against the planning horizon.

Because the results in Fig 5 did not include combinations of mitigation policies, Table 1 summarizes results over the 81 policy combinations evaluated. Three combinations emerged as optimal for some length of planning horizon. For a planning horizon of 1 to 15 years, the baseline policy (5% refuge, no localized ban, no areawide spray, no Bt seed tax) continued to be optimal even as resistance increased. For a planning horizon of 16 to 22 years, the optimal policy increased the refuge requirement from 5% to 20%, but did not impose a localized ban, an areawide spray, or Bt seed tax. For a planning horizon of 23 to 25 years, technically adding the 50% tax to the 20% refuge requirement was optimal, but the increase in economic surplus was trivial (<0.05%). Therefore, our economic surplus criterion suggested that the optimal resistance mitigation policy was no intervention if a shorter (≤15 years) planning horizon was used and, if a longer (≥16 years) planning horizon was used, increasing the required refuge to 20% for all farmers when resistance emerged. Because mitigation policies that increased the required refuge decreased current benefits to achieve increased future benefits, discounting implies that the 20% and 50% refuge policies would have generated less surplus than plotted in Fig 5. Calculations showed that, with a 13% or higher discount rate, the no-intervention baseline remained the optimal policy for all planning horizons less than or equal to 25 years.

**Table 1.**
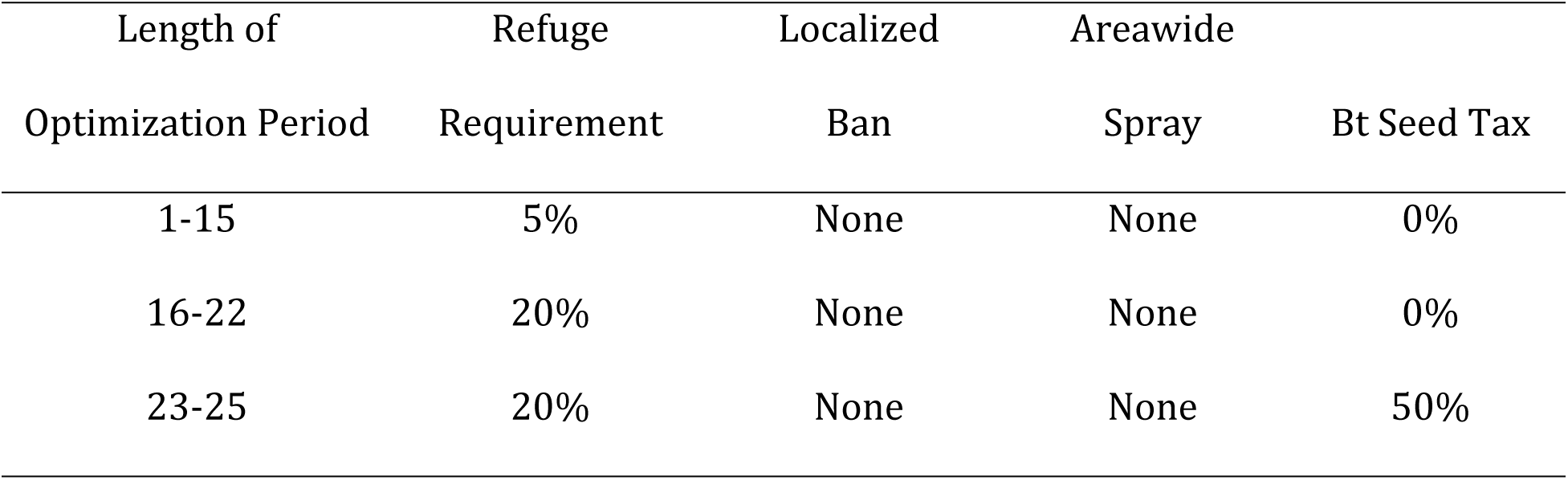
Optimal policy combination by length of optimization period

We also investigated the distribution of surplus shares under three resistance mitigation policies: the baseline (with a 5% refuge), a 20% refuge for all farmers planting Bt maize, and both a 20% refuge and a 50% Bt seed tax for all farmers planting Bt maize. Recall that economic surplus was the sum of farmer profit, the technology fee collected by the company and tax revenue and that adding the Bt seed tax as a mitigation policy had little impact on surplus with a 20% refuge requirement. For the baseline, farmers and the companies roughly divided the surplus evenly as yield gains and technology fees (Fig S1). Increasing the refuge requirement from the baseline of 5% to 20% to mitigate resistance increased the company share of surplus by about 5 to 10 percentage points, with the farmer share falling to about 40% (Fig S1). Adding a 50% Bt seed tax on top of the 20% refuge requirement to mitigate resistance, the tax burden was borne more by the companies, with their share declining by about 15 percentage points to 45% of the surplus, while farmers received about 35% of the surplus and tax revenue accounts for about 20% of the surplus.

### Role of Social Networks

To highlight the difference created by incorporating the effects of social networks, Fig 6 and Fig Fig 7 show results with all parameters the same as for the baseline except that the model was recalibrated with social networks “shut off” by setting the parameter *q* = 1. In this case, farmers made Bt maize adoption decisions based only on their individual expected profitably, giving no weight to their neighbors’ decisions. In terms of Bt maize adoption, without the effect of social networks, the farmer adoption rate grew faster first, but then slowed and eventually declined from period 43 onward (Fig 6). This result was explained by the lack of social network effects. Without them, profitable adoption by early adopters was not slowed by neighboring non-adopters. Similarly, as the technology became less effective due to resistance, profit-motivated dis-adoption of Bt maize was not slowed by neighbors’ inertia. As a consequence of the lower usage of Bt maize, the R-allele frequency reached key levels later than for the baseline. Specifically, the R-allele frequency did not exceed 10% until period 42, 20% until period 45, 30% until period 46, 40% until period 47, and 50% until period 48, or about 7 years later than for the baseline. Hence, not including the effects of social networks on farmer adoption of Bt maize slowed the estimated evolution of resistance by about 7 years.

**Fig 6.**
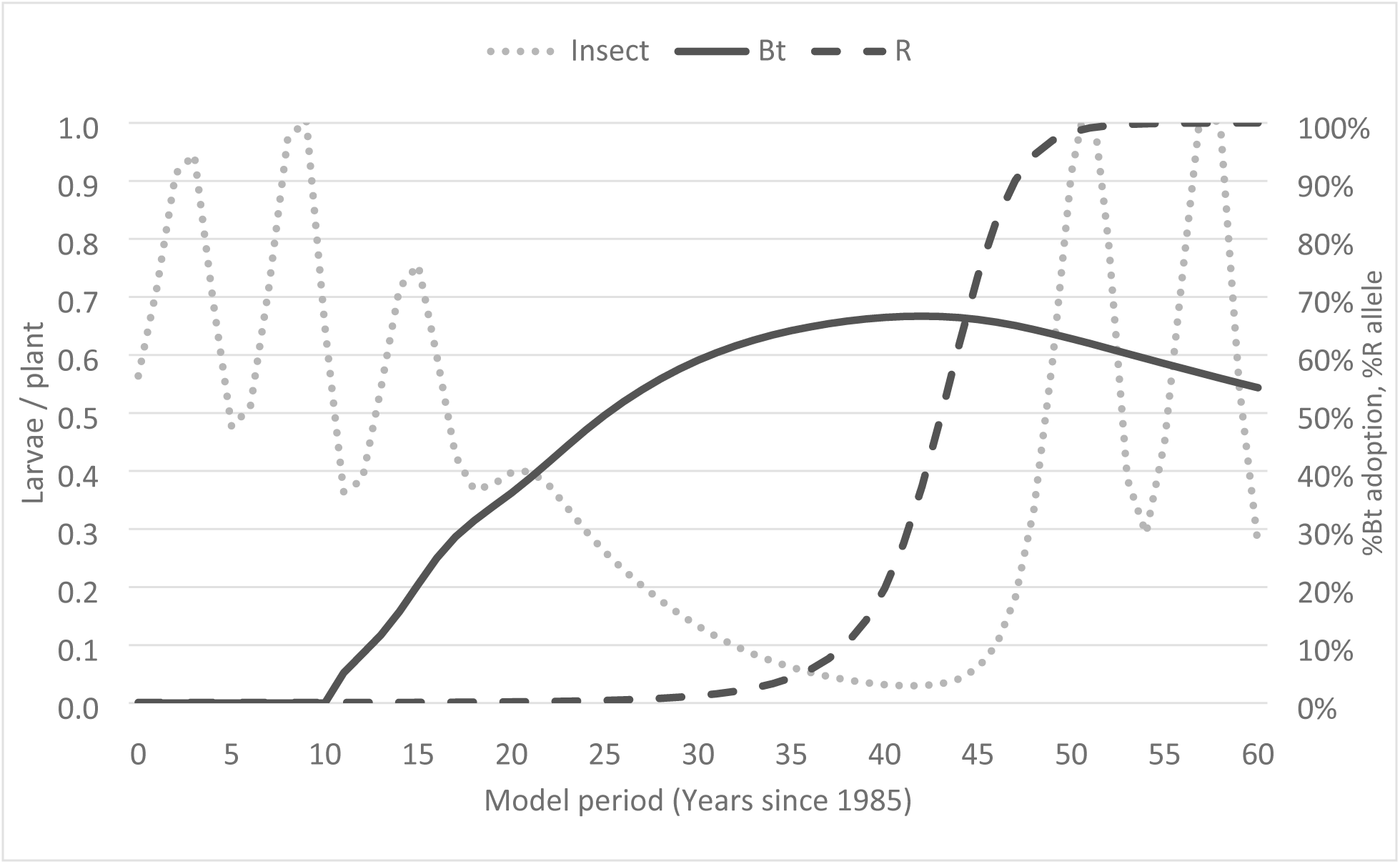
Results from the calibrated model without social network effects. It contains the insect population density (Insect), Bt seed adoption rate (Bt), and the resistance allele frequency (R) for the calibrated model without social network effects. The results for each period are averages over 1,000 simulations.

**Fig 7.**
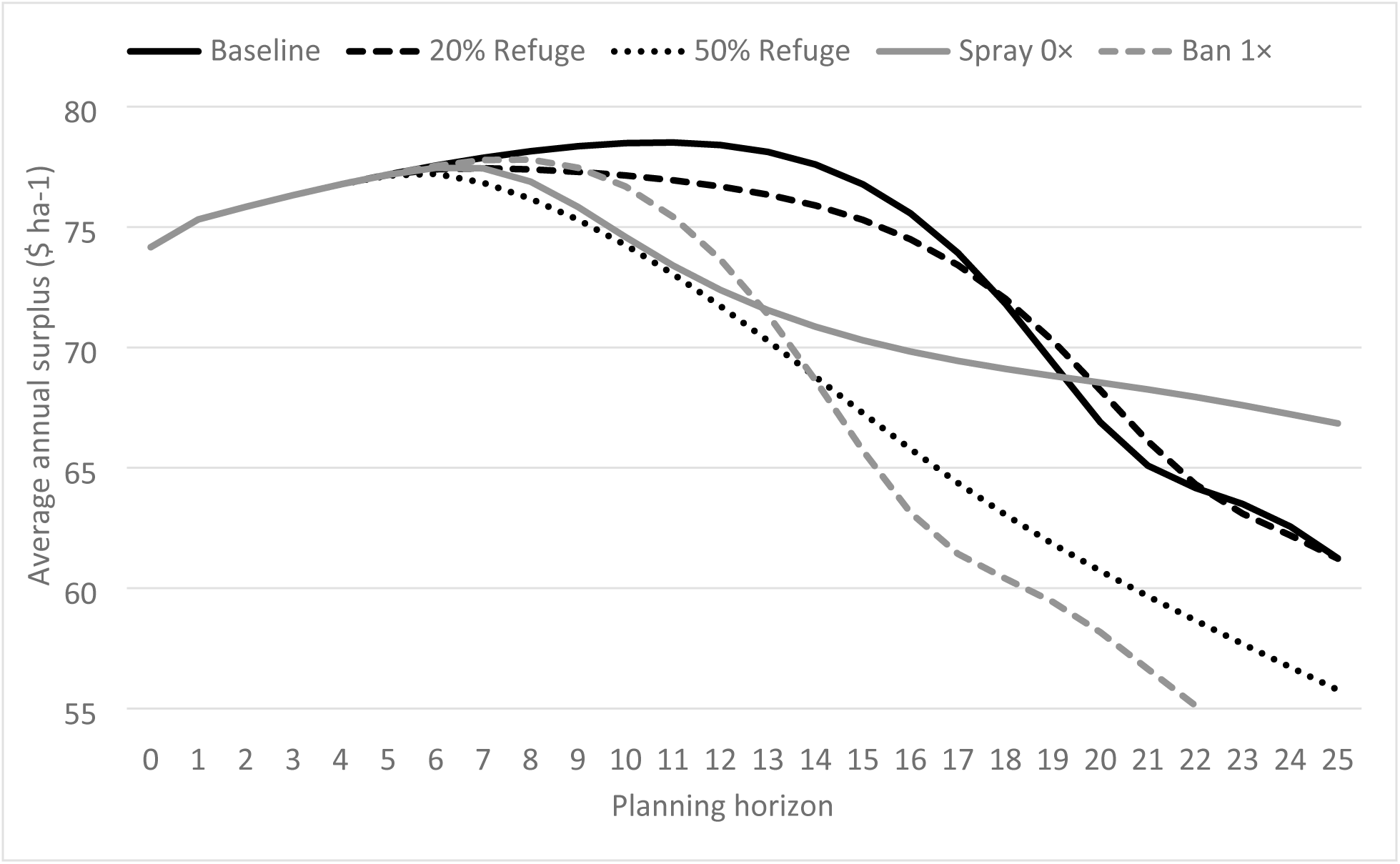
Average annual surplus for different mitigation policies. Each is plotted against the planning horizon for the calibrated model without social network effects.

Fig 7 plots average annual economic surplus against the planning horizon with the effect of social networks on adoption “shut off.” Again, results for the tax policies were omitted as they were almost identical to baseline results. Compared to Fig 5, which incorporated the effects of social networks on adoption, Fig 7 shows that all mitigation policies generated essentially the same surplus for the first 6 or 7 years. Because mitigation policies were not implemented until the R-allele frequency exceeded a 50% threshold, the slower projected evolution of resistance without social networks effects delayed policy implementation, so that all policies were initially equivalent.

Based on Fig 7, the optimal policy depended on the planning horizon and varied from Fig 5. The baseline again generated the greatest average annual surplus for all planning horizons up to 17 years (about the same as in Fig 5). Again, the 20% refuge for all farmers was the optimal mitigation policy for longer planning horizons, but only for the narrow range from 18 to 20 years. Furthermore, the difference between the baseline (with 5% refuge) and the 20% refuge mitigation policy was much smaller than in Fig 5. However, just as in Fig 5, the 50% refuge policy generated among the lowest amounts of economic surplus. Interestingly, for planning horizons exceeding 20 years, the areawide spray policy became optimal, which did not occur in Fig 5. This result occurred because without social network effects, farmers more quickly dis-adopted Bt maize when resistance developed, thus avoiding the higher costs of the spray policy and lower Bt maize benefits, and so they generated higher surplus. Again, the ban policy generated the lowest economic surplus over many planning horizons, even with the more rapid dis-adoption of Bt maize when resistance developed.

Because the results in Fig 7 did not include combinations of mitigation policies, Table 2 summarizes results over the 81 policy combinations evaluated, just as Table 1 did for Fig 5. Without social network effects, farmer adoption of Bt maize only responded to individual profitability, which created some shifts in the optimal mitigation policy. Three policy combinations again emerged as optimal for some length of planning horizon. For a planning horizon of 1 to 17 years, the baseline policy continued to be optimal even as resistance increased, and for a planning horizon of 18 to 19 years, the optimal policy increased the refuge requirement from 5% to 20%. These were the same policies as when the effects of social networks were included, but the planning horizons changed to be slightly longer for the baseline policy and shorter for the 20% refuge policy (Table 1). The greatest change without social network effects was for the longest planning horizons. For a 20- to 25-year planning horizon, the optimal resistance mitigation policy was to reduce the refuge requirement back to 5% for all farmers, to add a 50% Bt maize seed tax on all farmers, and to make areawide insecticide applications in areas where resistance emerged. With social network effects and longer planning horizons of 22 to 25 years, the refuge remained at 20% and only the 50% Bt seed tax was added (Table 1). The greater responsiveness of farmers to individual profitability without social network effects made more active mitigation policies optimal, but only for longer planning horizons. However, the effect was not large, as again calculations showed that with a 9% or higher discount rate, the no-intervention baseline remained the optimal policy for all planning horizons less than 25 years.

**Table 2.**
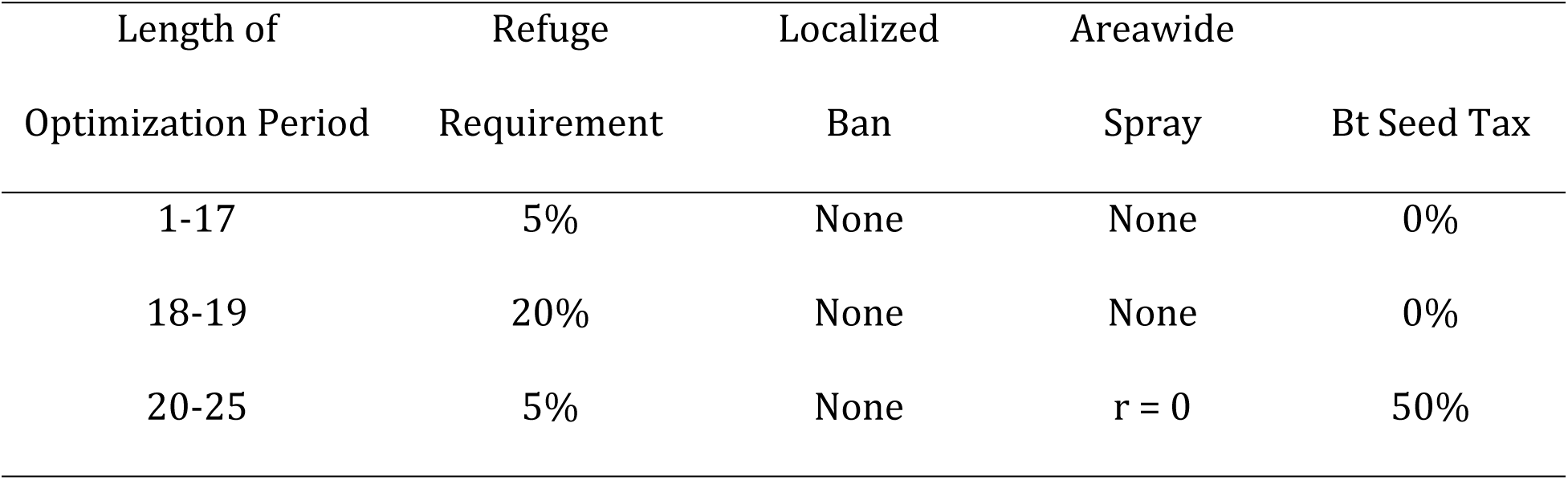
Optimal policy combination by length of optimization period without social network effects

## Discussion

Research on insect resistance mitigation strategies and empirical applications of agent-based models to pest and resistance management are limited [22–24]. Hence, as part of this paper’s first goal, we demonstrated the capacity of an agent-based model to produce results of use by policymakers and other stakeholders, specifically examining resistance mitigation policies for Bt maize and the European corn borer and the role of social networks in Bt maize adoption. We evaluated 81 resistance mitigation policies that combined three levels of four policies (non-Bt maize refuge, areawide non-Bt insecticide sprays, localized Bt maize bans, Bt maize seed tax) implemented when and/or where resistance emerged. These combinations showed variation in projected dynamics for Bt maize adoption, resistance allele frequency in the pest population, and the average pest population density.

From a biological perspective focused on keeping the frequency of resistance alleles in the pest population low, the most effective mitigation policies were a 50% refuge requirement for all farms when resistance emerged on the landscape and a localized ban on planting Bt maize within one radius of adult dispersal of farms with resistance. Bringing a broader economic perspective that balanced costs and benefits, we used economic surplus (the sum of farmer profit from maize production, company technology fees from selling Bt maize seed, and any tax revenue collected) to identify recommended mitigation policies. Surprisingly, results showed that when resistance emerged, the optimal response in terms of maximizing economic surplus was making no policy changes, but continuing the current resistance management policy of 5% non-Bt refuge, with no requirement of insecticidal sprays or localized Bt maize bans in and around areas of resistance, or Bt seed taxes when resistance emerges. For planning horizons beyond 16 years it became optimal to increase refuge requirements to 20% for all farmers when resistance developed. Furthermore, for planning horizons beyond 22 years it became optimal to add a 50% tax on all Bt maize seed sold when resistance developed in addition to the 20% refuge requirement. These results show the impacts that incorporating broader social science perspectives into resistance management or mitigation can have on recommended policy responses.

Several caveats apply to these results, as models cannot avoid the fundamental tradeoff between fidelity to the phenomenon examined and abstraction for ease of analysis and interpretation. Baseline results assume one single-toxin Bt maize producing a high dose of the toxin. However, multiple single-toxin Bt maize hybrids with different modes of action have been commercialized in the US, and single-toxin Bt maize hybrids have been phased out as companies have shifted to Bt maize hybrids with multiple, pyramided traits [35]. Furthermore, refuge requirements in the Midwest have changed over time for the different Bt maize hybrids. Initial requirements were for 20% non-Bt maize as structured refuge, but more recently, some Bt maize hybrids with pyramided traits have a 5% or 10% refuge requirement implemented as a seed mix and/or structured refuge [35, 36]. Our model does not capture the use of multiple toxins entering the market at different times, overlapping use of hybrids with multiple, pyramided traits at the same time by neighboring farmers, or changes in refuge requirements and methods of implementation. In addition, our model assumes that Bt maize delivers a high dose of the toxin, which is accurate for European corn borer, but not for other lepidopteran pests such as corn earworm (*Helicoverpa zea*) or Bt maize for corn rootworm [14,37,38]. Furthermore, the model focuses on a single pest, though farmers simultaneously manage multiple pests with varying levels of control by different Bt maize hybrids [39]. In addition, the model assumes a single selection by the Bt toxin each year, while many target pests, including the European corn borer, have multiple generations per season with more than one selection event by Bt maize [39]. Also, economic surplus is not a complete measure of social benefits [40]. For example, as used here, it does not include environmental impacts of insect management, even though a significant benefit of Bt maize is that farmers use it as a substitute for conventional insecticides [41–43].

With these caveats, the policy experiments reported here suggest that refuge requirements remain the foundation of resistance mitigation for high-dose technologies, just as they are for resistance management. Based on maximizing social surplus, the optimal policy to mitigate resistance when it emerges was to maintain the current refuge requirement or to modestly increase it for all farmers, rather than to implement localized bans on the sale of Bt maize in areas where resistance develops or to make areawide applications of insecticidal sprays on resistant populations. Based on the economic surplus criterion, the benefits of lower resistance allele frequency for these policies did not adequately compensate for the added costs or loss of the benefits from using Bt maize. Taxes on the sale of Bt maize seed did not cause surplus to differ substantially from the baseline policy, suggesting a possible mechanism to fund various programs to improve Bt maize use, such as development of educational materials and outreach or research activities. However, the results showed that companies bear a large share of these costs, suggesting that it would be more efficient for companies to directly fund these programs based on seed sales rather than creating a seed tax program to fund them. Also, as a caveat, this model did not incorporate the Bt technology market. As a result, for example, the model did not include market competition among companies via differentiated traits, including different regulatory requirements, as, for example, companies would lobby to not have their hybrids included in tax schemes if resistance developed to a competitor’s Bt maize.

As a secondary goal of this paper, we demonstrated that social factors can play key roles in the development and management of insect resistance, focusing on the effect that social networks can play on farmer adoption of Bt maize. As modeled here, adoption depended in part on the average adoption of a farmer’s neighbors, not just each farmer’s expected profitability, as a way to capture the effects of information exchange, integration of multiple farmers’ experiences with pests and adoption, shared local institutions and markets, and similar factors. Modeling the mechanisms for this social network and the specific connections among individual farmers is beyond the scope of this analysis. Relative to a model in which farmers responded only to their individual profitability, social networks as modeled here impeded farmer responsiveness to profitability signals, which slowed the initial adoption of Bt maize and its dis-adoption as pest populations declined or resistance developed. Model calibration to observed state-level adoption rates identified model parameters and reduced differences in initial adoption rates with and without social networks. However, this calibrated model implied a relatively slower adjustment in Bt maize use by farmers. As a result, when including social network effects, our model projected that resistance develops about 7 years earlier than without social network effects and the optimal mitigation policy more strongly favored use of moderate increases in refuge for all farmers. Resistance developed earlier because farmers uses Bt maize more intensely since they did not dis-adopt Bt maize as pest populations declined and resistance developed, even though the profitability of Bt maize decreased. With social network effects, the optimal resistance mitigation policy also more strongly favored use of modest increases in refuge because more farmers continued to use Bt maize and obtain it benefits relative to more costly, but effective, policies such as areawide sprays or localized bans. In this example, ignoring social network effects could contribute to making inappropriate policy recommendations for managing pest resistance or mitigating it once it develops.

The intensity and extent of farmer adoption of Bt maize plays a key role in the management and mitigation of pest resistance. This agent-based model incorporated the influence of social factors by having individual farmer adoption respond to expected profitability and the adoption behavior of neighboring farmers. However, many social factors not addressed by this model also affect adoption. For example, expected profitability depends not only on all the market factors driving maize prices, but also technology markets and the pricing behavior of firms selling Bt maize [29]. Similarly, farmers adopt Bt maize not only for expected profit, but also to manage income risk [44, 45]. Also, social networks for agricultural management rarely have the simple spatial structure assumed here, but typically have varying nodes of importance such as key crop consultants, retailers, and extension agents [30, 46]. In addition, social factors affect resistance through more than just adoption of Bt maize, such as through farmer compliance with resistance management and mitigation practices and how Bt maize affects broader cropping systems such as crop rotations [10, 11]. Overall, our results demonstrate that social factors can play an important role in resistance management and mitigation. However, more applied and quantitative social science research would contribute to developing better policy recommendations for resistance management and mitigation, and agent-based models can be a part of this contribution.

## Model

### Landscape

The spatially explicit model used a 30×70 grid space representing the cropland in Wisconsin. Modeled farmers mimic the Wisconsin crop landscape [47] and plant 44% of the fields to maize, the host crop for the pest. Fields maintain their initial random assignment to maize or non-maize production during a simulation, but are reassigned at the start of each new simulation. During a simulation, maize farmers decide adoption of Bt maize each period. Fig 8 depicts a typical model landscape, in this case with 90% Bt adoption and a resistance allele frequency of 51% for the total population. A circle (**○**) represents a farmer who plants conventional (non-Bt) maize, whereas a black dot (●) represents a farmer who adopts Bt maize. A light-gray background (■□) indicates that the pest population in an individual field before adult dispersal has a resistance allele frequency of more than 50%, the criterion used for declaring that a population is resistant [48]. To avoid boundary effects, top fields wrap to corresponding bottom fields and left-most fields to corresponding right-most fields, creating a torus, implying that the model space is part of a larger landscape with comparable dynamics occurring for the pest population and its genetic structure [28].

**Fig 8.**
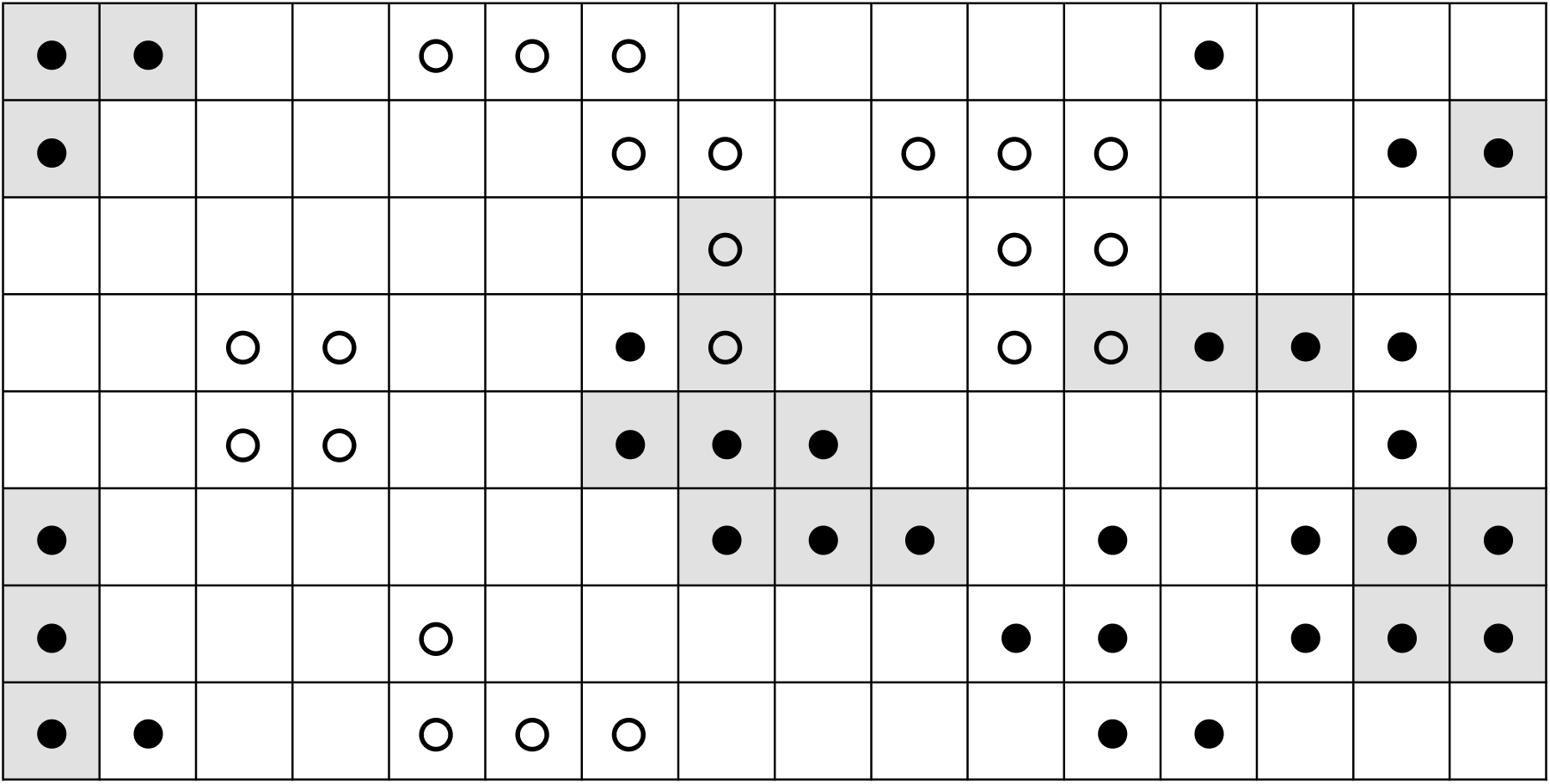
Example model landscape. Circle (○) represents a field planted to non-Bt maize, black dot (●) represents a field planted to Bt maize, and light-gray (■□) indicates a field with a pest population with a resistance allele frequency of more than 50% before adult dispersal.

### Pest Population Genetics

The pest population-genetics model is parametrized for European corn borer (*Ostrinia nubilalis*), a major pest for Midwestern maize and the primary target for initial commercial releases of Bt maize beginning in 1996 [32]. The insect model uses discrete time steps corresponding to distinct generations and consistent with the seasonality of many types of crop production and pest life cycles. *O. nubilalis* typically has two generations per year in the major US maize production region, though northern regions may have only one generation per year and southern regions may have three or more [39]. The model simplifies these dynamics to one discrete time step per year that aggregates population dynamics and genetic selection across these generations. Hutchison et al. [32] used a comparable empirical approach to estimate annual population growth rates for *O. nubilalis* using annual observations of second-generation adult population densities in Minnesota and Wisconsin.

Historically, the *O. nubilalis* population in the Midwestern US has oscillated with an approximately seven year cycle [49] largely due to the entomopathogenic parasite *Nosema pyrausta* [31]. Field data for second-generation populations in Wisconsin over 1944-1995 show an average peak and trough for the oscillation of about 1.2 and 0.2 larvae per plant [31, 32]. The population model approximates these dynamics using a lagged logistic growth model:

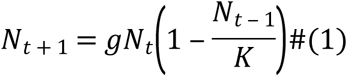

where *N_t_* is the second-generation larval population (larvae per maize plant), *g* is the annual growth rate and *K* is the carrying capacity. Using *g* = 2.15 and *K* = 1.4 generates a reasonable approximation of historical *O. nubilalis* population dynamics in Wisconsin, with a similar range of population minimums and maximums as observed and six or seven years between peaks.

The genetics model assumes two alleles, *R* for resistant and *S* for susceptible, creating three genotypes, homozygous resistant *RR*, homozygous susceptible *SS*, and heterozygous *RS*, with respective Bt toxin larval survival rates of 1.0, 0.0, and 0.18. Each period, after Bt toxin mortality and density-dependent mortality for larvae, random mating occurs among the adult population within each field before adult dispersal. Note that, with random mating, *RR* and *RS* genotypes both contribute *R* alleles, but *RS* genotypes do so half as often on average, also contributing *S* alleles just as often. Let *α_t_*, *β_t_*, and *γ_t_* respectively denote the relative frequencies in period *t* of *RR*, *SS*, and *RS* genotypes, which by definition sum to 1. Random mating then implies 1 = *α_t_* _+ 1_ + *β_t_* _+ 1_ + *γ_t_* _+ 1_ = (*α_t_* + *β_t_* + *γ_t_*)^2^ = (*α_t_* + 0.5*γ_t_*)^2^ + (*β_t_* + 0.5*γ_t_*)^2^ + 2(*α_t_* + 0.5*γ_t_*)(*β_t_* + 0.5*γ_t_*), so that

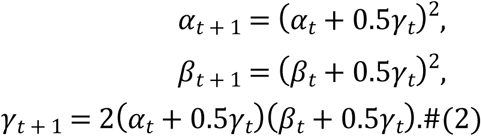

Adults disperse uniformly within a radius *r* of maize fields (i.e. not onto non-maize fields). The literature provides a range of observations for dispersal, with dispersal in most cases taking place within 20km and depending on various factors (season, gender, mating status). To capture the effects of dispersal yet remain computationally tractable, the model uses a dispersal radius of 15km, which corresponds to 3 fields in the model grid space. We assume that the natal field is always available as a destination, which implies that if no neighboring fields exist within the dispersal range, adults stay in the same field.

### Farmer Behavior

Individual farmers manage each field and decide each period whether to plant Bt or non-Bt maize. A number of economic and social factors influence farmers’ adoption decisions for Bt maize [50]. Rather than explicitly enumerating and modeling these multiple factors, agent-based models combine simple behavioral models with suitable random components and let complex phenomenon emerge [18]. Though expected profitability greatly influences farmer management decisions in commercial agriculture, their local social networks also significantly influence their behaviors, not just by providing additional information regarding the relative profitability of different practices [30,46,51]. Therefore, we model the Bt adoption process as a hybrid of individual profit maximization and local imitation to capture the effect of social networks.

### Farmer Profit

The profit-based component of farmer behavior uses the following switching function [25]:

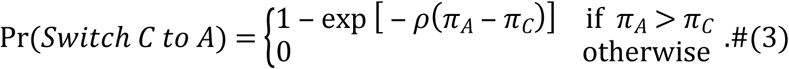

Here, *π_A_* is the profitability of the alternative choice and *π_C_* is the profitability of the current choice, with both profits calculated using the pest population density for the previous period in the field. The function determines the probability that the farmer switches from the current choice to the alternative (*Switch C to A*), with the probability increasing as the alternative becomes relatively more profitable than the current choice. We use a “soft” probabilistic switching decision to capture the effect of other unobserved individual factors [39]. The parameter *ρ* captures the responsiveness of farmer adoption to profitability differences, with a greater *ρ* increasing the probability farmers use the more profitable alternative. As explained in the Calibration section, we calibrate *ρ* against the Bt seed adoption data for Wisconsin to derive *ρ* = 0.0023. The negative-exponential function implies that the switching probability is the farmers’ expected utility gain from switching when the gain is uncertain, assuming constant absolute risk aversion for the farmer, a commonly used assumption for empirical analysis [52, 53].

Farmer profit for a field (*π*) is crop revenue minus cost, where revenue declines as the pest population increases and cost varies with the scenario:

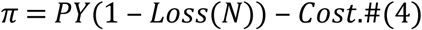

Here *P* is crop price ($ Mg^-1^), *Y* is potential or pest-free crop yield (Mg ha^-1^), *Loss* is proportional crop loss, which depends on *N*, the average pest population density (larvae per plant), and *Cost* is the production cost ($ ha^-1^). To focus on factors other than annual variability in crop prices and yields, crop price and potential yield are fixed at reported averages in 2017 for Wisconsin farmers: *P* = $129.91 ha^-1^ and *Y* = 10.92 Mg ha^-1^ [54]. These values imply constant potential revenue of $1,418.62 ha^-1^ across fields and seasons. The proportion of potential revenue lost due to pest damage depends on the average larval population density based on an empirical model [44]: *Loss*(*N*) = 0.1186*N*^0.5146^.

*Cost* consists of a base cost *C* ($ ha^-1^) that does not vary by policy scenario and costs that do:

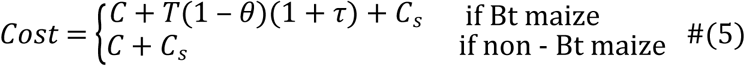

Based on US Department of Agriculture crop budgets [55], the base cost *C* is set to $1,202.51 ha^-1^, the reported average for 2017 in the region containing Wisconsin for all costs except opportunity costs for land and operator labor and management. *T* ($ ha^-1^) is the additional seed cost for Bt maize (“technology fee”), which varies over time based on the function estimated with Wisconsin market data [27]. Specially, *T* = $17.49 ha^-1^ from 1996 to 2003, and then declines to *T* = $17.45 ha^-1^ for 2004, $15.78 ha^-1^ for 2006, $13.75 ha^-1^ for 2007, $11.41 ha^-1^ for 2008, $9.18 ha^-1^ for 2009, $8.29 ha^-1^ for 2010, $7.82 ha^-1^ for 2011, $7.39 ha^-1^ for 2012, and then remains at a base of $7.04 ha^-1^ for years 2013 and afterward. The remaining cost parameters vary with the policy scenario: *θ* is the proportion of refuge (non- Bt maize) planted with Bt maize, *τ* is the tax rate for Bt maize, and *C_s_* is the cost ($ ha^-1^) for a foliar insecticidal spray as part of areawide management of adults. Each scenario sets these cost parameters at appropriate values as described in the Policy Experiments section. For example, a refuge only scenario sets *τ* = *C_s_* = 0 and sets *θ* at 0.05, 0.20 or 0.50; a ban only scenario sets *τ* = *C_s_* = 0 and *θ* = 1 (100% refuge) in fields where a ban is in effect; and an areawide spray policy sets *τ* = 0 and imposes the cost *C_s_* for all affected fields. The cost of an insecticidal spray *C_s_* is $33.51 ha^-1^ based on published survey averages for insecticide active ingredients used in maize and application costs, adjusted for inflation to 2017 equivalents [33, 34].

### Social Network

Network analysis has been widely applied to understand the diffusion of innovations as a social phenomenon, including in agriculture [56, 57]. Neighboring farmers have been shown to create a local environment that affects individual farmer adoption decisions, both for hybrid maize seed and for Bt maize [30, 58]. To capture this social network effect, the model assumes each farmer in a field is connected to farmers in neighboring fields, with the size of the neighborhood determined by a “radius”. Fig 9 shows an example of a size-2 neighborhood for a farmer with nine neighbors who plant maize, either Bt or non-Bt. Those neighbors themselves have their own neighborhoods, with each connection undirected so that the local social networks are tightly overlapped. The number of neighbors for a size-*n* neighborhood can range from 0 to a maximum of 4*n*(*n* + 1). With no data for social network sizes for farmers, and considering that *n* = 3 gives up to 48 neighbors (implying a substantial computational burden), the model randomly assigns neighborhood sizes to each farmer for all seasons using a uniform distribution over {0, 1, 2}.

**Fig 9.**
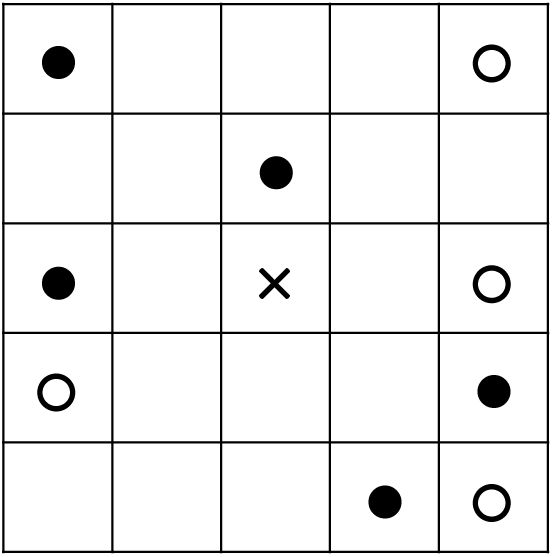
Example size-2 neighborhood. It is centered on a farmer (×) with nine neighbors who plant maize, either Bt maize (●) or non-Bt maize (○).

Given this local social network, each maize farmer chooses each season to grow either Bt or non-Bt maize for a field. A parameter *q* defines the impact of social networks on farmer adoption decisions. With probability *q*, a farmer focuses solely on individual profits using the switching function and with probability 1 ‒ *q* follows the majority choice of his neighbors in the previous season. For example, if the farmer in Fig 9 follows the majority, he plants Bt maize next season because his neighborhood has 5 Bt maize adopters and 4 non-adopters. In the case of a tie, he chooses Bt maize as well. Also, one-third of the farmers randomly have a size-0 neighborhood (*n* = 0) and so, with no social network, always use the switching function. As a result, the probability that a farmer uses the switching function is 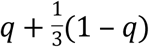, while the probability that a farmer has a size-1 neighborhood is 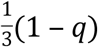, which is also the probability that a farmer has a size-2 neighborhood. Thus, the model has two calibration parameters for Bt maize adoption: *q* defining the impact of social networks and *ρ* defining the responsiveness of farmers to profit in the switching function.

Lastly, a refuge policy is implemented as a fixed proportion *θ* of non-Bt maize with complete compliance by farmers, the so-called “refuge in a bag” [59] in which the company mixes Bt and non-Bt maize seeds before purchase. In our model, the refuge requirement has two effects. First, the effective seed cost is the proportion (1 ‒ *θ*) of the technology fee *T* at that period. Second, the effective survival rate of each genotype is calculated as the weighted average: *θ* + (1 ‒ *θ*)*s*, where *s* is the original survival rate. That is, with probability *θ*, any genotype survives due to the non-Bt maize, and with probability 1 ‒ *θ*, each genotype survives according to its Bt toxin survival rate. Initially, we assume *θ* = 0.05, which is the lowest refuge requirement already in place, and later increased refuge levels are examined as resistance mitigation policies.

### Running the Model

Each model run begins with initialization, including randomly placing farmers across the landscape. Since corn fields occupy roughly 44% of total farmland in Wisconsin (represented by 30×70 fields configured as a torus), the total number of maize farmers for a run is approximately 0.44 × 30 × 70 = 924. After initialization, the run proceeds period by period, with a period corresponding to a growing season or year. Before introducing Bt maize into the model, the insect module runs for 11 periods, which corresponds to the pre-Bt periods and helps stabilize the model’s biological dynamics. Thereafter, the model simultaneously updates the pest population density of each field for each period. First, Bt toxin effects reduce each field’s pest population based on the survival rates of the genotypes established there the previous season. Second, mating determines the genotype composition of the next generation based on random mating of the population in the field. Third, reproduction determines the pest population density based on the lagged logistic growth model. Fourth, the population locally re-mixes across fields based on the dispersal model. Finally, maize farmers simultaneously make planting decisions (whether to plant Bt or non-Bt maize) for the next period based on the farmer behavioral model. In short, during a growing season the Bt toxin (if present) reduces the natal population in a field, survivors randomly mate and produce the next generation, which then disperses locally across fields, and then farmers make maize planting decisions for the following spring.

### Calibration

We used aggregate Bt maize adoption data for Wisconsin to calibrate the model. Our calibration minimized the average of the mean squared error (MSE) of prediction for the simulated landscape compared to the observed data. Specifically, the MSE for a run was the squared deviation of the simulated Bt adoption rate from the annual Wisconsin adoption data, averaged across all periods with adoption data (*t* = 11 to 32). Since runs were random, the MSEs were averaged across 1,000 runs. The two calibration parameters were the responsiveness of farmers to expected profit differences between alternatives (*ρ*) and the probability (*q*) that farmers focus solely on profit differences to make adoption decisions, rather than their neighbors’ choices. To avoiding both over-fitting and excessive computational requirements, a grid search was used with increments of 0.0002 for *ρ* and of 0.1 for *q*. To highlight the significance of local networks, we also calibrated the model by fixing *q* = 1 and using only *ρ*, which “shut off” all social network effects on adoption.

Plotting the Wisconsin Bt maize adoption data and both calibration fits shows the superior fitting of the two-parameter hybrid model relative to the one-parameter model (Fig 10). Using the same random seeds for both models, the optimum solutions are *ρ* = 0.0036 and *q* = 0.3 for the hybrid model and *ρ* = 0.0022 for the single parameter model. These optimal values for the two-parameter model imply that 70% of the years, farmers follow the majority choice of their neighborhood, suggesting that network effects are important for understanding farmer adoption dynamics for Bt maize.

**Fig 10.**
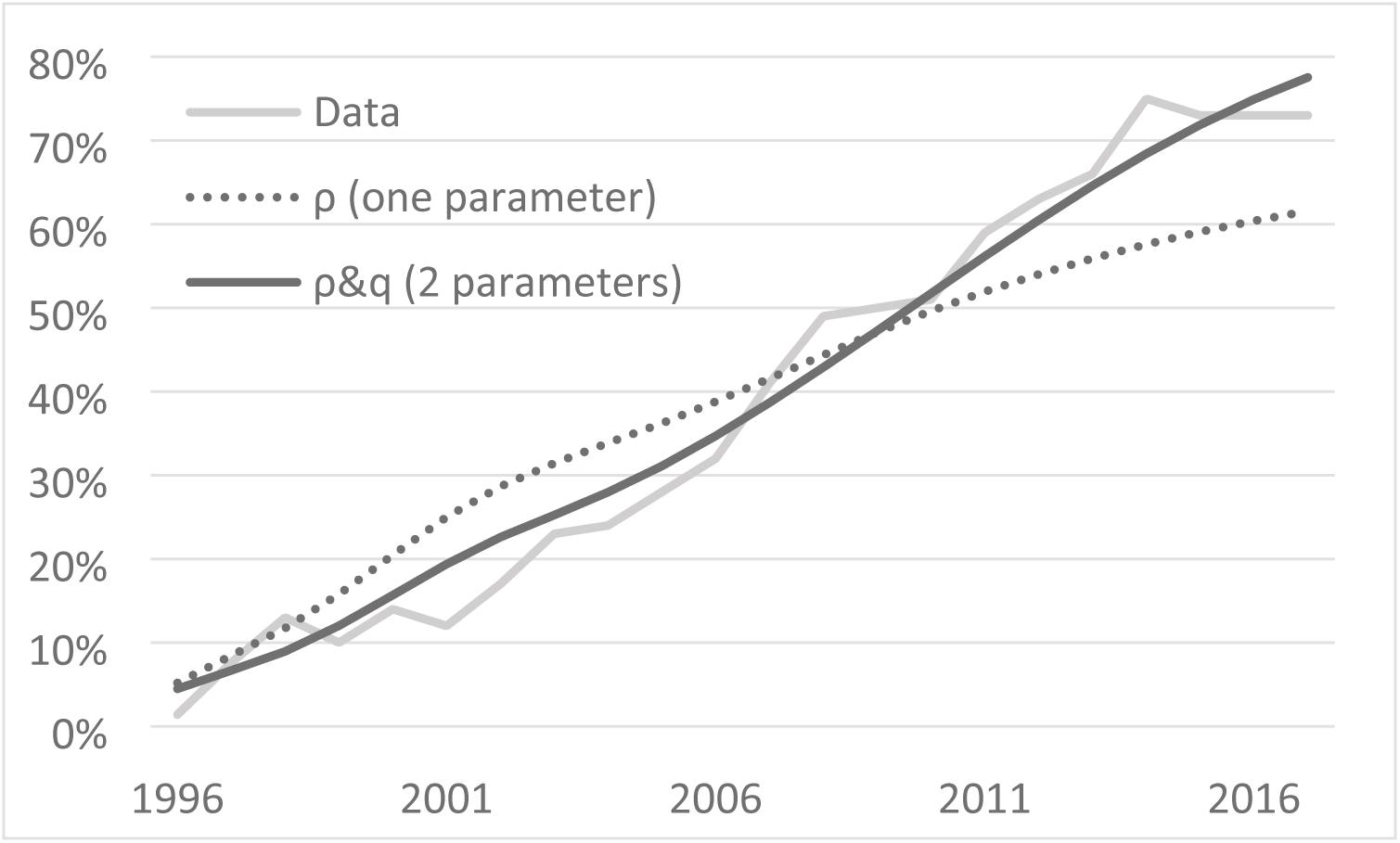
Aggregate adoption of Bt maize in Wisconsin and two simulated results. The simulated results are generated by calibrating one parameter and two-parameters.

## Acknowledgments

We wish to acknowledge the financial support of the U.S Department of Agriculture National Institute for Food and Agriculture’s Agriculture and Food Research Initiative (grant number 2014-67023-21814 awarded to TMH and PDM, https://nifa.usda.gov/program/agriculture-and-food-research-initiative-afri) and Monsanto’s Corn Rootworm Knowledge Research Program (grant awarded to PDM and TMH: https://monsanto.com/products/product-stewardship/corn-rootworm-management/corn-rootworm-knowledge-research-program/. The funders had no role in study design, data collection and analysis, decision to publish, or preparation of the manuscript.

## Supporting information

**Fig S1.**
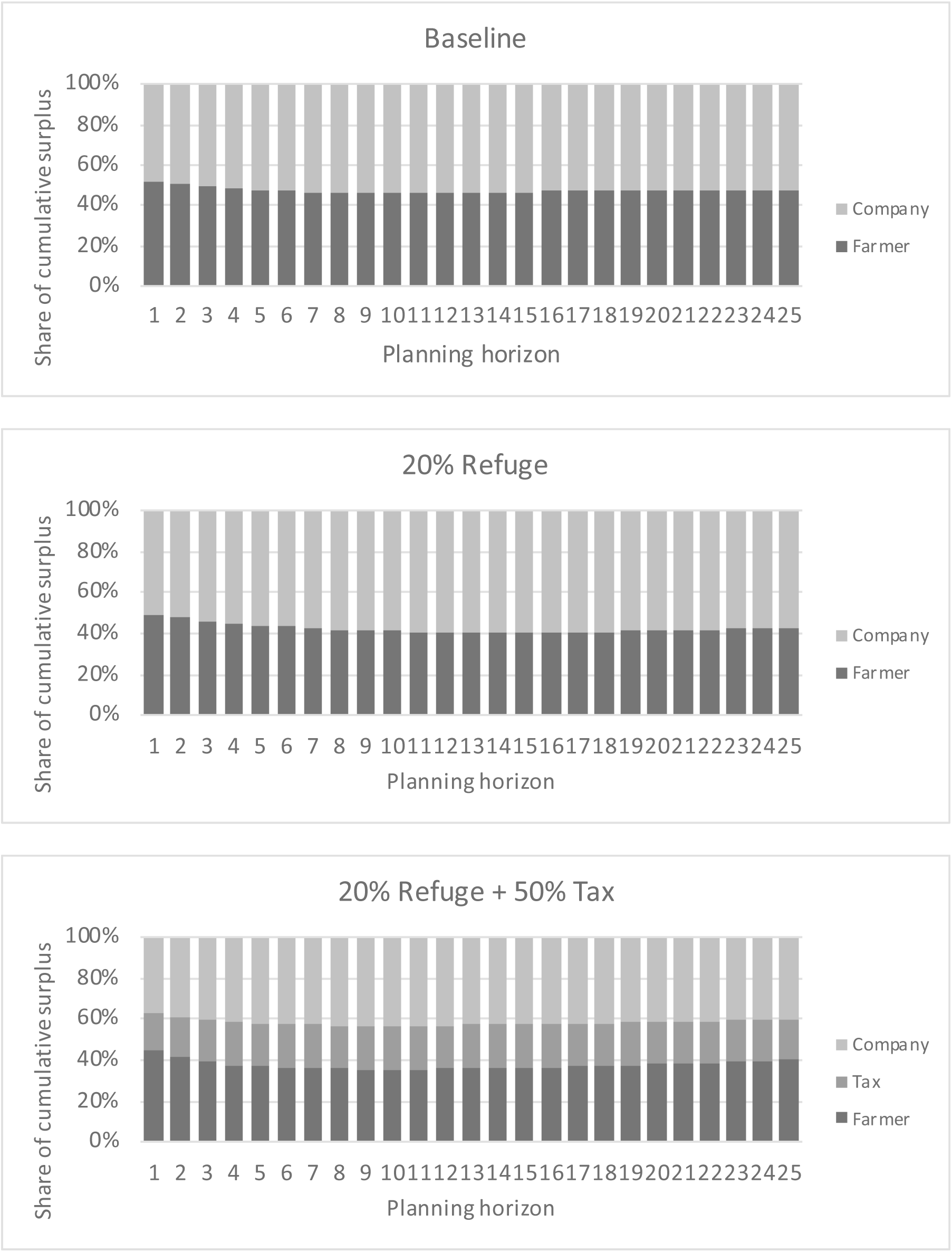
Cumulative share of surplus by planning horizon. The top panel is for the baseline, the middle is for the 20% refuge mitigation policy, and the bottom is for the 20% refuge mitigation policy combined with a 50% Bt seed tax.

**Fig S2.**
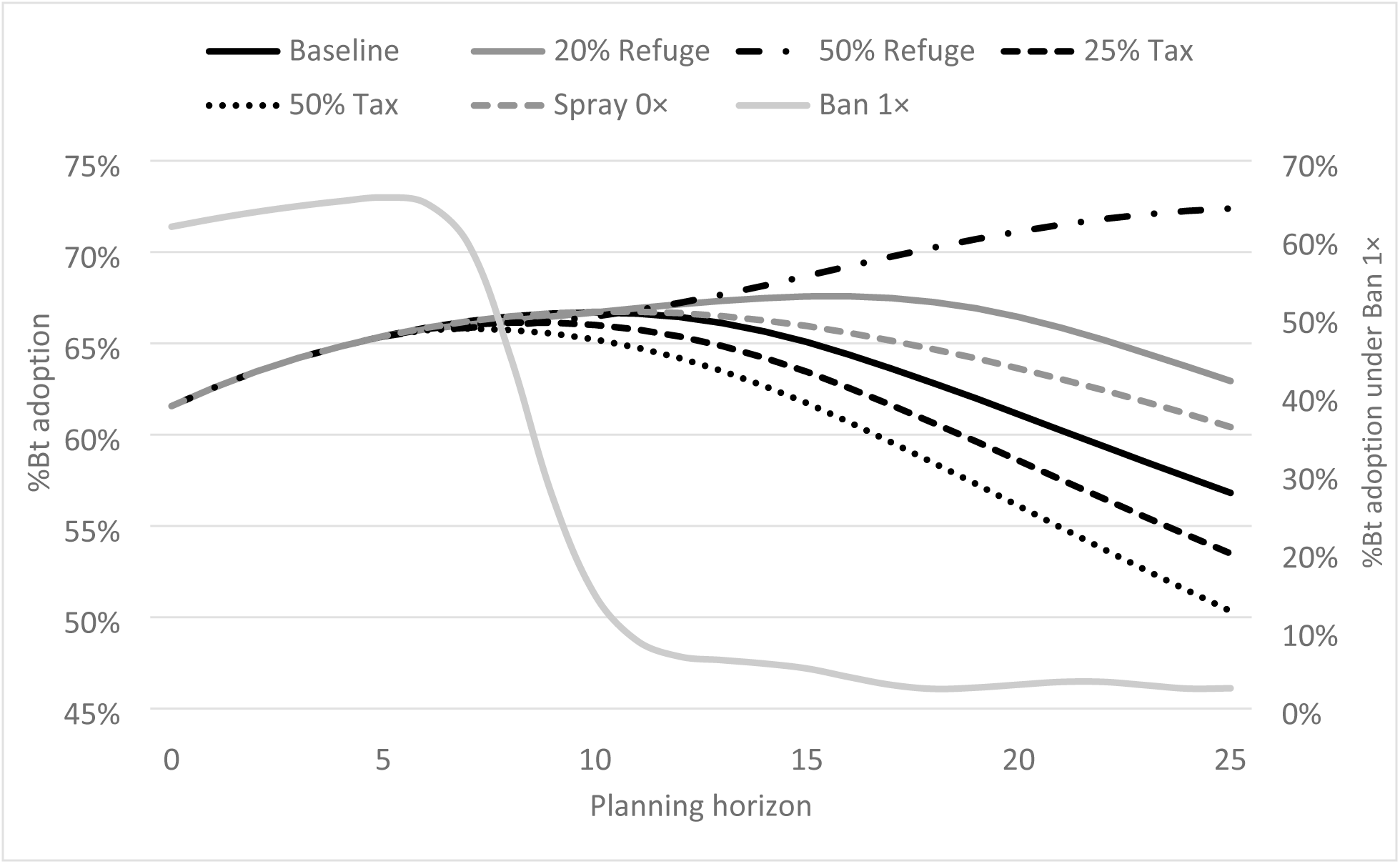
Bt adoption rate under simple policies. Each is plotted against the planning horizon without social network effects. The results for each period are averages over 1,000 simulations.

**Fig S3.**
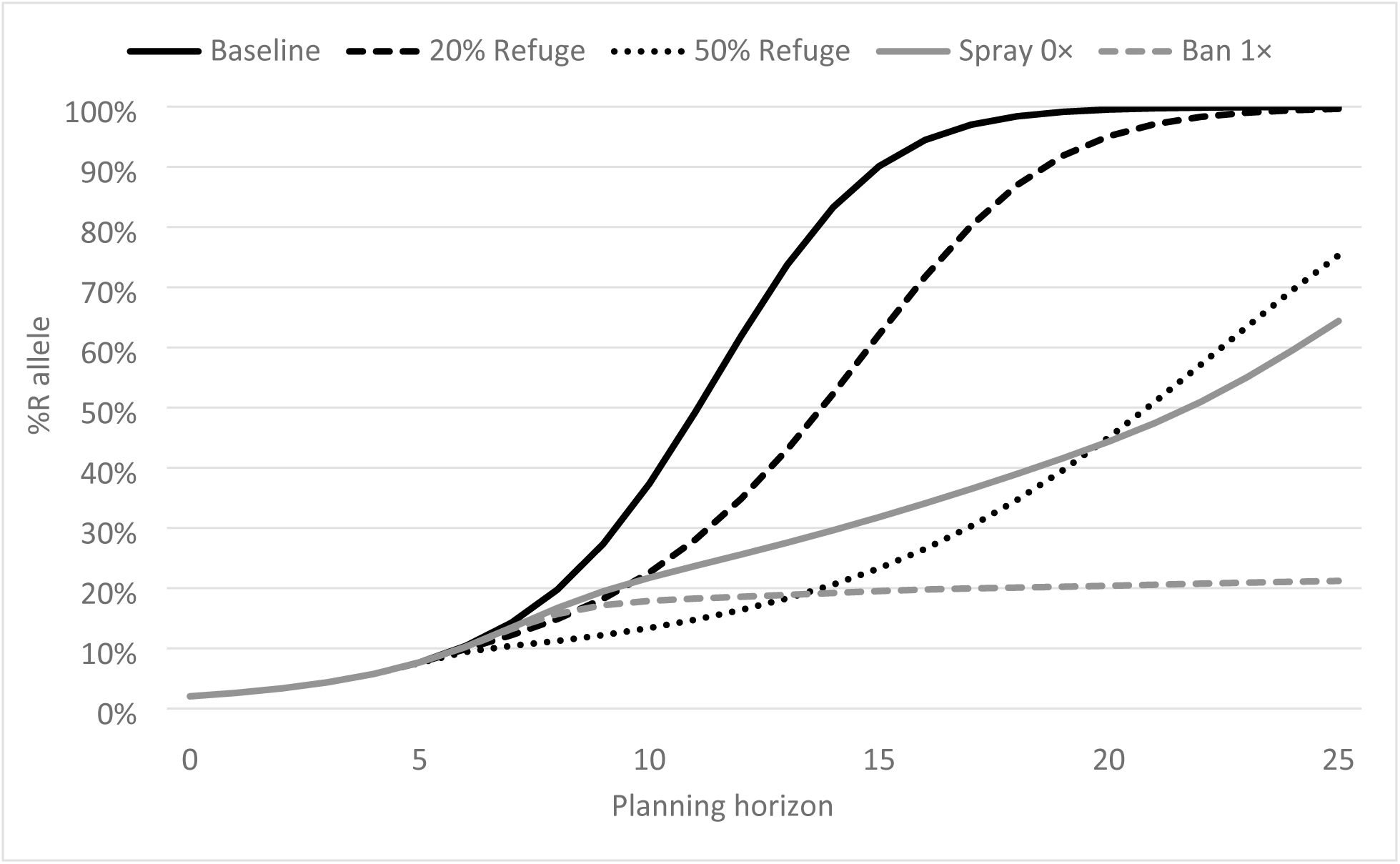
R-allele frequency under simple policies. Each is plotted against the planning horizon without social network effects. The results for each period are averages over 1,000 simulations.

**Fig S4.**
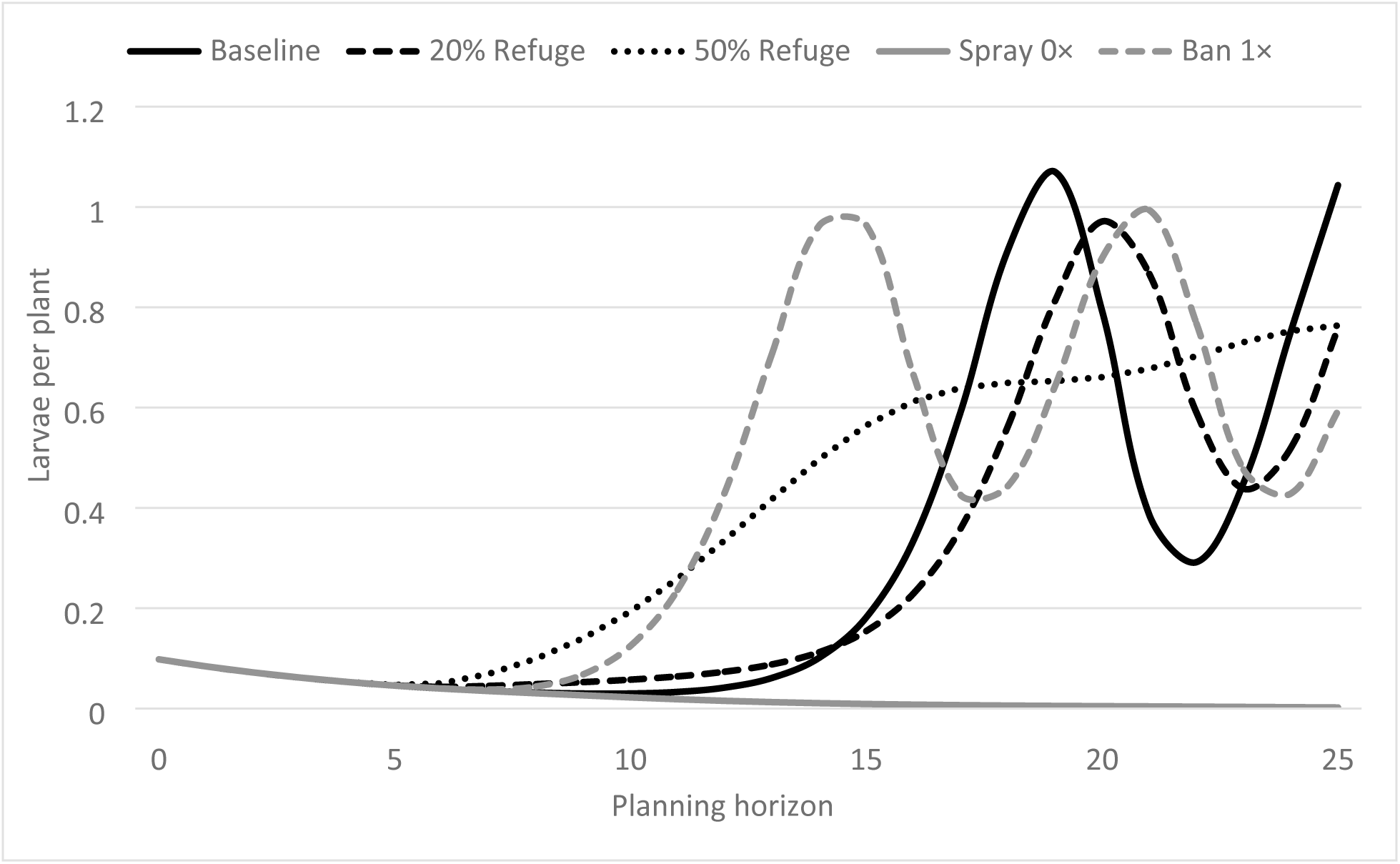
Insect population density under simple policies. Each is plotted against the planning horizon without social network effects. The results for each period are averages over 1,000 simulations.

## References

1. ISAAA. Global Status of Commercialized Biotech/GM Crops in 2017: Biotech Crop Adoption Surges as Economic Benefits Accumulate in 22 Years. Ithaca, NY: ISAAA; 2017. (ISAAA Brief). Report No.: 53.

2. USDA. World Agricultural Supply and Demand Estimates [Internet]. Washington, DC; 2018. (WASDE). Report No.: 581. Available from: https://www.usda.gov/oce/commodity/wasde/

3. Tabashnik BE. ABCs of Insect Resistance to Bt. PLOS Genet. 2015 Nov 19;11(11):e1005646.

4. Huang F, Andow DA, Buschman LL. Success of the high-dose/refuge resistance management strategy after 15 years of Bt crop use in North America. Entomol Exp Appl. 2011 Jul 1;140(1):1–16.

5. Pardo-López L, Soberón M, Bravo A. Bacillus thuringiensis insecticidal three-domain Cry toxins: mode of action, insect resistance and consequences for crop protection. FEMS Microbiol Rev. 2013 Jan 1;37(1):3–22.

6. Tabashnik BE, Brévault T, Carrière Y. Insect resistance to Bt crops: lessons from the first billion acres. Nat Biotechnol. 2013 Jun;31(6):510–21.

7. Gassmann AJ, Petzold-Maxwell JL, Keweshan RS, Dunbar MW. Field-Evolved Resistance to Bt Maize by Western Corn Rootworm. PLOS ONE. 2011 Jul 29;6(7):e22629.

8. Hurley TM, Babcock BA, Hellmich RL. Bt corn and insect resistance: an economic assessment of refuges. J Agric Resour Econ. 2001;26(1):176.

9. Crowder D, Onstad D, Gray M, Mitchell P, Spencer J, Brazee R. Economic analysis of dynamic management strategies utilizing transgenic corn for control of western corn rootworm (Coleoptera: Chrysomelidae). J Econ Entomol. 2005;98(3):961–75.

10. Mitchell PD, Onstad DW. Effect of extended diapause on evolution of resistance to transgenic Bacillus thuringiensis corn by northern corn rootworm (Coleoptera: Chrysomelidae). J Econ Entomol. 2005;98(6):2220–2234.

11. Hurley TM, Mitchell PD. Insect Resistance Management: Adoption and Compliance. In: Onstad DW, editor. Insect Resistance Management. 2nd ed. San Diego: Academic Press; 2014. p. 421–51.

12. Rebaudo F, Dangles O. An agent-based modeling framework for integrated pest management dissemination programs. Environ Model Softw. 2013 Jul 1;45:141–9.

13. USDA. Biopesticides Regrisrtaiotn Action Dcouments: Cry1Ab and Cry1F Bacillus thuringiensis (Bt) Corn Plant-Incorporated Protectants. Washington, DC; 2010.

14. Gassmann AJ, Petzold-Maxwell JL, Clifton EH, Dunbar MW, Hoffmann AM, Ingber DA, et al. Field-evolved resistance by western corn rootworm to multiple Bacillus thuringiensis toxins in transgenic maize. Proc Natl Acad Sci. 2014;201317179.

15. Andow DA, Pueppke SG, Schaafsma AW, Gassmann AJ, Sappington TW, Meinke LJ, et al. Early Detection and Mitigation of Resistance to Bt Maize by Western Corn Rootworm (Coleoptera: Chrysomelidae). J Econ Entomol. 2016 Feb 1;109(1):1–12.

16. Miller JH, Page SE. Complex adaptive systems: An introduction to computational models of social life. Princeton university press; 2009.

17. Peck SL. Simulation as experiment: a philosophical reassessment for biological modeling. Trends Ecol Evol. 2004 Oct 1;19(10):530–4.

18. Epstein JM. Generative social science: Studies in agent-based computational modeling. Princeton University Press; 2006.

19. Durlauf SN. Complexity, economics, and public policy. Polit Philos Econ. 2012 Feb 1;11(1):45–75.

20. Filatova T, Verburg PH, Parker DC, Stannard CA. Spatial agent-based models for socio- ecological systems: Challenges and prospects. Environ Model Softw. 2013 Jul 1;45:1–7.

21. Feola G, Binder CR. Towards an improved understanding of farmers’ behaviour: The integrative agent-centred (IAC) framework. Ecol Econ. 2010 Oct 15;69(12):2323–33.

22. Renton M. Shifting focus from the population to the individual as a way forward in understanding, predicting and managing the complexities of evolution of resistance to pesticides. Pest Manag Sci. 2013;69(2):171–5.

23. Renton M, Busi R, Neve P, Thornby D, Vila-Aiub M. Herbicide resistance modelling: past, present and future. Pest Manag Sci. 2014;70(9):1394–404.

24. Gay P-E, Lecoq M, Piou C. Improving preventive locust management: insights from a multi-agent model. Pest Manag Sci. 2017;74(1):46–58.

25. Milne AE, Bell JR, Hutchison WD, van den Bosch F, Mitchell PD, Crowder D, et al. The effect of farmers’ decisions on pest control with Bt crops: a billion dollar game of strategy. PLOS Comput Biol. 2015;11(12).

26. Ives AR, Paull C, Hulthen A, Downes S, Andow DA, Haygood R, et al. Spatio-Temporal Variation in Landscape Composition May Speed Resistance Evolution of Pests to Bt Crops. PLOS ONE. 2017 Jan 3;12(1):e0169167.

27. Onstad DW, Meinke LJ. Modeling Evolution of Diabrotica Virgifera Virgifera (Coleoptera: Chrysomelidae) to Transgenic Corn with Two Insecticidal Traits. J Econ Entomol. 2010 May 31;103(3):849–60.

28. Storer NP. A spatially explicit model simulating western corn rootworm (Coleoptera: Chrysomelidae) adaptation to insect-resistant maize. J Econ Entomol. 2003;96(5):1530–1547.

29. Shi G, Chavas J, Stiegert K. An Analysis of the Pricing of Traits in the U.S. Corn Seed Market. Am J Agric Econ. 2010 Oct 1;92(5):1324–38.

30. Kaup BZ. The reflexive producer: The influence of farmer knowledge upon the use of Bt corn. Rural Sociol. 2008;73(1):62.

31. Bell JR, Burkness EC, Milne AE, Onstad DW, Abrahamson M, Hamilton KL, et al. Putting the brakes on a cycle: bottom-up effects damp cycle amplitude: Managing epidemic persistence. Ecol Lett. 2012 Apr;15(4):310–8.

32. Hutchison WD, Burkness EC, Mitchell PD, Moon RD, Leslie TW, Fleischer SJ, et al. Areawide Suppression of European Corn Borer with Bt Maize Reaps Savings to Non-Bt Maize Growers. Science. 2010 Oct 8;330(6001):222–5.

33. Mitchell PD. Methods and assumptions for estimating the impact of pyrethroid insecticides on pest management practices and costs for U.S. crop farmers [Internet]. Madison, WI: AgInfomatics; 2017. Available from: http://aginfomatics.com/pyrethroids-project.html

34. USDA. Agricultural Prices [Internet]. 2018. Available from: https://usda.library.cornell.edu/concern/publications/c821gj76b

35. Hurley TM, Sun H. Softening Shock and Awe Pest Management in Corn and Soybean Production with IPM Principles. J Integr Pest Manag. 2019;10(1):7.

36. U.S. Environmental Protection Agency. Insect Resistance Management for Bt Plant-Incorporated Protectants [Internet]. 2019 [cited 2019 Jun 13]. Available from: https://www.epa.gov/regulation-biotechnology-under-tsca-and-fifra/insect-resistance-management-bt-plant-incorporated

37. U.S. Environmental Protection Agency. White Paper on Resistance in Lepidopteran Pests of Bacillus thuringiensis (Bt) Plant-Incorporated Protectants in the United States [Internet]. 2018. Available from: https://www.epa.gov/sites/production/files/2018-07/documents/position_paper_07132018.pdf

38. Burkness EC, Dively G, Patton T, Morey AC, Hutchison WD. Novel Vip3A Bacillus thuringiensis (Bt) maize approaches high-dose efficacy against Helicoverpa zea (Lepidoptera: Noctuidae) under field conditions: Implications for resistance management. GM Crops. 2010;1(5):337–43.

39. Mason CE, Rice ME, Calvin DD, Van Duyn JW, Showers WB, Hutchison WD, et al. European Corn Borer Ecology and Management. Iowa State University; 1996. (North Central Regional Extension Publication). Report No.: 327.

40. Mitchell PD, Brown Z, McRoberts N. Economic issues to consider for gene drives. J Responsible Innov. 2018;5(sup1):S180–202.

41. Perry ED, Ciliberto F, Hennessy DA, Moschini G. Genetically engineered crops and pesticide use in US maize and soybeans. Sci Adv. 2016;2(8):e1600850.

42. Catarino R, Areal F, Park J, Parisey N. Spatially explicit economic effects of non- susceptible pests’ invasion on Bt maize. Agric Syst. 2019;175:22–33.

43. Brookes G. Twenty-one years of using insect resistant (GM) maize in Spain and Portugal: farm-level economic and environmental contributions. GM Crops Food. 2019;1–12.

44. Hurley TM, Mitchell PD, Rice ME. Risk and the value of Bt corn. Am J Agric Econ. 2004;86(2):345–358.

45. Shi G, Chavas J-P, Lauer J. Commercialized transgenic traits, maize productivity and yield risk. Nat Biotechnol. 2013;31(2):111.

46. Lubell M, Fulton A. Local policy networks and agricultural watershed management. J Public Adm Res Theory. 2007;18(4):673–96.

47. USDA. Census of agriculture [Internet]. 2012. Available from: https://www.agcensus.usda.gov/Publications/2012/

48. Tabashnik BE. Delaying insect resistance to transgenic crops. Proc Natl Acad Sci. 2008 Dec 9;105(49):19029–30.

49. Onstad DW, Maddox JV. Modeling the effects of the microsporidium, Nosema pyrausta, on the population dynamics of the insect, Ostrinia nubilalis). J Invertebr Pathol. 1989;53(3):410–21.

50. Useche P, Barham BL, Foltz JD. Integrating Technology Traits and Producer Heterogeneity: A Mixed-Multinomial Model of Genetically Modified Corn Adoption. Am J Agric Econ. 2009 May 1;91(2):444–61.

51. McAllister R, Robinson C, Maclean K, Guerrero A, Collins K, Taylor B, et al. From local to central: a network analysis of who manages plant pest and disease outbreaks across scales. Ecol Soc. 2015;20(1).

52. Chavas J-P. Risk analysis in theory and practice. Academic Press; 2004.

53. Mitchell PD, Hutchison WD. Decision making and economic risk in IPM. In: Radcliffe EB, Cancelado RE, Hutchison WD, editors. Integrated Pest Management: Concepts, Tactics, Strategies and Case Studies [Internet]. Cambridge: Cambridge University Press; 2008. p. 33–50. Available from: https://www.cambridge.org/core/books/integrated-pest-management/decision-making-and-economic-risk-in-ipm/1B17130CD5A60148EA1057C0B1ECE4AF

54. USDA. Quick Stats Lite [Internet]. 2019. Available from: https://www.nass.usda.gov/Quick_Stats/Lite/

55. USDA. Commodity Costs and Returns [Internet]. 2016 [cited 2016 Oct 28]. Available from: http://www.ers.usda.gov/data-products/commodity-costs-and-returns/commodity-costs-and-returns/#Recent Costs and Returns: Corn

56. Easley D, Kleinberg J. Networks, crowds, and markets: Reasoning about a highly connected world. Cambridge University Press; 2010.

57. Jackson MO. Social and economic networks. Vol. 3. Princeton university press; 2010.

58. Ryan B, Gross NC. The diffusion of hybrid seed corn in two Iowa communities. Rural Sociol. 1943;8(1):15.

59. Hodgson E. Refuge in a Bag is Here: Explaining the Simplified Refuge [Internet]. 2010 [cited 2018 Jul 17]. Available from: https://crops.extension.iastate.edu/cropnews/2010/06/refuge-bag-here-explaining-simplified-refuge

